# Mechanosensing by the lamina protects against nuclear rupture, DNA damage, and cell cycle arrest

**DOI:** 10.1101/583179

**Authors:** Sangkyun Cho, Manasvita Vashisth, Amal Abbas, Stephanie Majkut, Kenneth Vogel, Yuntao Xia, Irena L. Ivanovska, Jerome Irianto, Manorama Tewari, Kuangzheng Zhu, Elisia D. Tichy, Foteini Mourkioti, Hsin-Yao Tang, Roger A. Greenberg, Benjamin L. Prosser, Dennis E. Discher

## Abstract

Whether cell forces or extracellular matrix (ECM) can impact genome integrity is largely unclear. Here, acute perturbations (~1hr) to actomyosin stress or ECM elasticity cause rapid and reversible changes in lamin-A, DNA damage, and cell cycle. Embryonic hearts, differentiated iPS-cells, and various nonmuscle cell types all show that actomyosin-driven nuclear rupture causes cytoplasmic mis-localization of DNA repair factors and excess DNA damage. Binucleation and micronuclei increase as telomeres shorten, which all favor cell cycle arrest. Deficiencies in lamin-A and repair factors exacerbate these effects, but lamin-A-associated defects are rescued by repair factor overexpression and by contractility modulators in clinical trials. Contractile cells on stiff ECM normally exhibit low phosphorylation and slow degradation of lamin-A by matrix-metalloprotease-2 (MMP2), and inhibition of this lamin-A turnover and also actomyosin contractility is seen to minimize DNA damage. Lamin-A is thus stress-stabilized to mechano-protect the genome.

## Introduction

Proliferation of many cell types slows dramatically shortly after birth and is absent in key adult tissues (Li, et al., 1996), which presents a major challenge to regeneration. Cell growth in culture is modulated by the stiffness of extracellular matrix (ECM) which generally promotes actomyosin stress (Engler, et al., 2006; Paszek, et al., 2005) and affects the conformation (Sawada, et al., 2006), post-translational modification (PTM) (Guilluy, et al., 2014), localization (Dupont, et al., 2011), and degradation (Dingal and Discher, 2014) of mechanosensitive proteins. Actomyosin links to ECM and to nuclei, such that rapid changes to ECM and tissue mechanics that result from chronic or acute injury or even drug therapies (e.g. cardiac arrest or cardiomyopathy treatment (Green, et al., 2016)) can in principle affect the nucleus (Takaki, et al., 2017; Wiggan, et al., 2017; Kanellos, et al., 2015) and perhaps the DNA within. DNA damage and telomere shortening are indeed well-documented in injuries that affect heart (Chang, et al., 2018; Higo, et al., 2017; Sharifi-Sanjani, et al., 2017; Oh, et al., 2003) as well as nonmuscle tissue – but DNA damage and repair are rarely studied in developing organs, and relationships to proliferation, ECM stiffness, and actomyosin stress are understudied.

DNA damage and senescence increase with many disease-linked mutations, including those in the nucleoskeletal protein lamin-A (LMNA) that forms a structural meshwork around chromatin (Turgay, et al., 2017; Shimi, et al., 2015; Gruenbaum, et al., 2005). *LMNA* deficiencies associate with elevated DNA damage (Graziano, et al., 2018; Liu, et al., 2005) and result in accelerated aging of stiff tissues similar to deficiencies in DNA repair factors (e.g. KU80) (Li, et al., 2007). Moreover, progeroid syndromes are caused only by mutations in *LMNA* and DNA repair factors, but LMNA’s primary function in development remains hotly debated (Burke and Stewart, 2013), with suggested roles in gene positioning and regulation (Harr, et al., 2015) seeming at odds with largely normal development of human and mouse mutants until weeks after birth. Surprisingly, senescence or apoptosis of cells with LMNA defects is rescued by culturing cells on almost any ECM (versus rigid plastic (de La Rosa, et al., 2013; Hernandez, et al., 2010)) and by treatment with at least one drug affecting both cytoskeleton and nucleo-cytoplasmic trafficking (Larrieu, et al., 2018; Larrieu, et al., 2014). Relationships between lamins, actomyosin stress, ECM mechanics, and DNA damage are nonetheless obscure – especially in tissues.

Embryonic hearts beat spontaneously for days after isolation from early chick embryos, and beating is acutely sensitive to myosin-II inhibition (**Fig.1A**) as well as enzymatic stiffening or softening of ECM (Majkut, et al., 2013). The latter studies reveal an optimal stiffness for beating that is likewise evident for cardiomyocytes (CMs) cultured on gels (Majkut, et al., 2013; Engler, et al., 2008; Jacot, et al., 2008). DNA damage is conceivably optimized in heart as it triggers a switch from proliferation to senescence in post-natal hearts (Puente, et al., 2014). DNA damage is also implicated in telomere attrition and binucleation of CMs that signal irreversible exit from cell cycle (Aix, et al., 2016). We postulated embryonic hearts with rapidly tunable mechanics could prove useful as a tissue model for clarifying protein-level mechanosensing mechanisms *in vivo* that could be studied thoroughly *in vitro* with many cell types.

**Figure 1.**
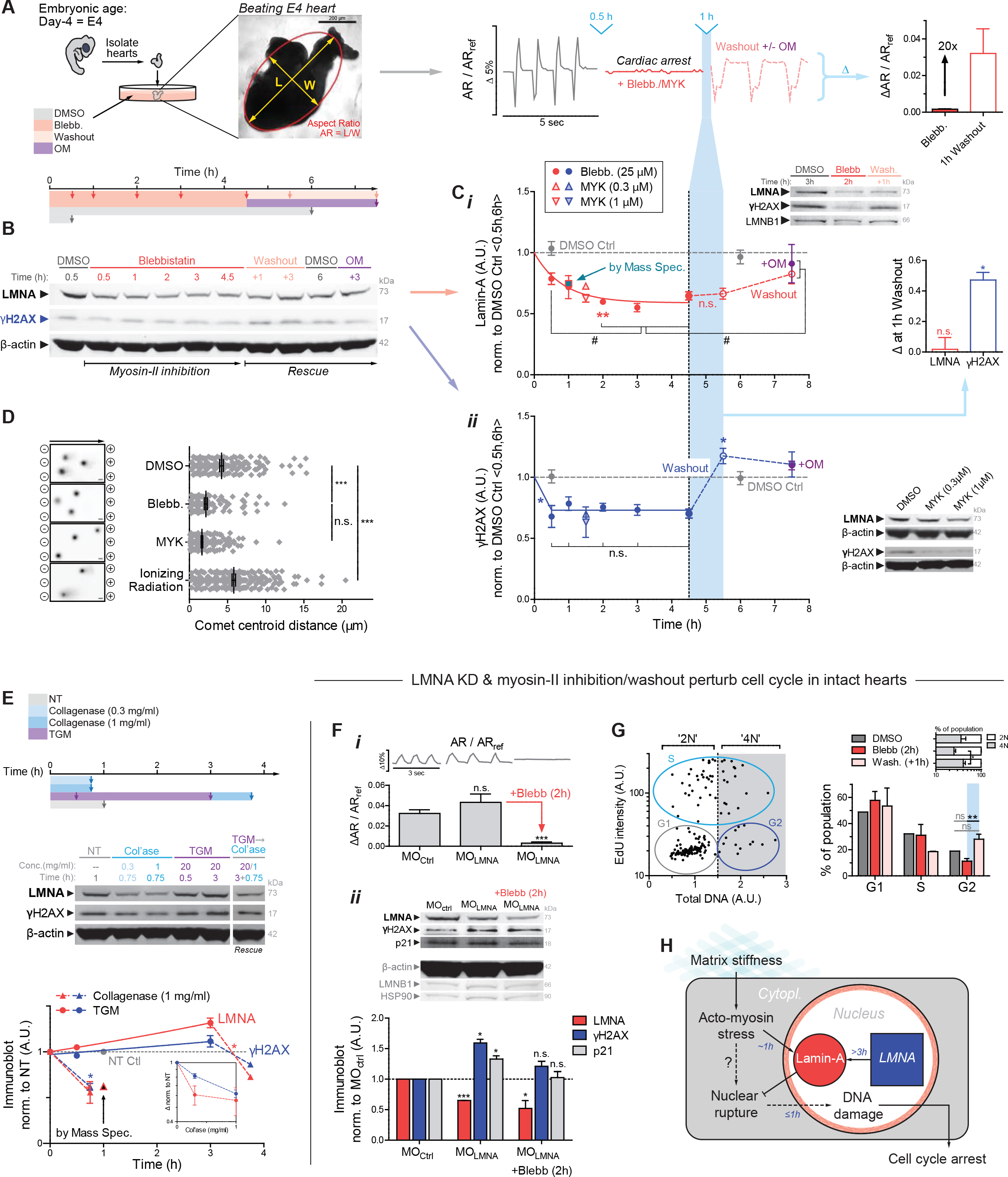
Contractility or collagen perturbations result in rapid ~1h changes in LMNA, DNA damage, and cell cycle. **(A)** Chick hearts from day 4 (E4) beat at 1-2 Hz for up to 5 d. Middle: Aspect ratio (AR) beating strain is arrested by myosin-II inhibition, but recovers with drug washout ± myosin-II activator, OM. **(B)** Immunoblot of hearts inhibited for varying durations, followed by washout ± OM (8 hearts per lysate). **(C) (*i*)** Immunoblot densitometry reveals LMNA decreases in ~45 min upon myosin-II inhibition, but LMNA recovery takes >3h upon washout. MS confirms 1h data with 29 LMNA-specific peptides, and inset shows LMNB1 remains unchanged. **(*ii*)** DNA damage γH2AX also decreases, but spikes upon blebb washout (5.5h, light blue). Bar graph of LMNA and γH2AX changes 1h post-washout. Bottom right immunoblot shows MYK treatment for 1.5 h also affects LMNA and γH2AX. One-way ANOVA (Dunnett’s test vs DMSO Ctrl avg of 0.5h,6h): \**p*<0.05, **<0.01; t-test (vs plateau average of 2, 3, 4.5h time points): #*p*<0.05. All error bars indicate ±SEM. **(D)** Electrophoretic Comet assay of DNA damage validates γH2AX quantitation, with γ-irradiation control. **(E)** Immunoblot of hearts treated with collagenase and/or transglutaminase (TGM) (8 hearts per lysate). Matrix digestion rapidly decreases LMNA and γH2AX, consistent with low stress; cross-linking has opposite and reversible effect by 3h (t-test: \**p*<0.05; superfluous blot lanes have been removed and indicated by a blank space). **(F) (*i*)** Morpholino KD of LMNA (MO_LMNA_) does not affect beating versus MO_Ctrl_, but **(*ii*)** MO_LMNA_ increases DNA damage (γH2AX) and cell cycle inhibitor p21, unless rescued by myosin-II inhibition (*i*,*ii*). Two immunoblots (anti-LMNA, p21, β-actin) & (anti-LMNB1, γH2AX, HSP90) display proteins of interest above loading controls. (8 hearts per lysate). One-way ANOVA: \**p*<0.05, ***<0.001. **(G) (*i*)** Representative plot of EdU vs DNA intensity for cell cycle phase (G1 vs S vs G2; or ‘2N’ vs ‘4N’). (*ii*)Blebb washout increases G2 and 4N cells (vs 2N). n>77 cells per cond.; t-test: \*\**p*<0.01. **(H)** Proposed mechanosensing pathway.

Although LMNA is reportedly ‘undetectable’ in early embryos (Stewart and Burke, 1987), it progressively accumulates in tissues such as heart, bone, and lung (Solovei, et al., 2013; Rober, et al., 1989; Lehner, et al., 1987) and is highest in adults within these mechanically stressed, stiff tissues that are collagen-rich (Swift, et al., 2013). LMNA deficiencies accordingly produce measurable defects weeks after birth in such tissues, including heart (Kubben, et al., 2011; Worman and Bonne, 2007). We therefore hypothesized that LMNA in normal embryos mechanosenses the earliest microenvironment in the first organ and increases not only to stiffen the nucleus (Osmanagic-Myers, et al., 2015; Dahl, et al., 2008; Pajerowski, et al., 2007) but also to regulate repair factors that confer resistance to both DNA damage and cell cycle arrest.

## Results

### Contractility and collagen perturbations rapidly impact LMNA and DNA damage in hearts

To first assess the consequences of cytoskeletal stress on nuclei in a tissue, we arrested embryonic hearts (day-4: E4) with reversible inhibitors of myosin-II (Fig.1A). Spontaneous beating of hearts stopped in minutes with blebbistatin (‘blebb’), but the heart re-started just as quickly upon washout of drug (**Fig.1A** **right**). An activator of cardiac myosin-II omecamtiv mecarbil (‘OM’) – in clinical trials for heart failure (NCT02929329) – re-started the hearts reliably. Quantitative immunoblots and mass spectrometry (MS) measurements (**Fig.1B**, **1C-*i***, **S1A&B**) revealed a rapid decrease in LMNA (~45 min decay constant) while LMNB remained unchanged (**Fig.1C-i**, **upper right inset**, **Fig.S1C**), and similar results were evident for a cardiac myosin-II-specific inhibitor ‘MYK’ (mavacamten analog) – in clinical trials for hypertrophic cardiomyopathy (NCT02842242). Despite rapid recovery of beating after drug washout (**Fig.S1D**), LMNA recovered more slowly (>3h), indicating slow synthesis.

DNA damage in post-natal development with LMNA deficiencies (Liu, et al., 2005) led us to hypothesize acute, embryonic changes in phospho-histone γH2AX, as a primary marker of DNA damage. A rapid *decrease* in DNA damage was surprising with myosin-II inhibition (**Fig.1C-*ii***) given the decrease LMNA, but electrophoretic comet assay confirmed the γH2AX results (**Fig.1D**). It is useful to keep in mind that the heart beats reasonably well with LMNA deficiencies and mutations. Because blebbistatin washout recovers beating while LMNA remains low, we anticipated a large spike in DNA damage shortly after washout (Fig.1C-*i*&*ii*, right inset). LMNA and DNA damage eventually reached control levels (in ~hrs), but the spike highlights the disruptive effects of actomyosin stress on genome integrity.

Actomyosin contractility is generally downstream of ECM stiffness (Ulrich, et al., 2009; Engler, et al., 2006), including for immature cardiomyocytes (CMs) (Engler, et al., 2008; Jacot, et al., 2008). Acute perturbations of collagen matrix might therefore be expected to affect DNA damage. Collagenase treatment for 45 min indeed resulted in rapid decreases in DNA damage and LMNA (**Fig.1E**), consistent with rapid softening of E4 hearts (~50%) and weaker beating (Majkut, et al., 2013). Treatment with tissue transglutaminase (TGM), a cross-linker of ECM that stiffens heart and thereby increases basal tension (>2-fold in 2h (Majkut, et al., 2013)), increased γH2AX and LMNA (only after 3h) except when collagenase was subsequently added (Fig.1E). LMNA thus decreases quickly or increases slowly in response to changes in ECM stiffness or actomyosin tension, both of which appear to also affect DNA damage. Effects are also generally reversible.

### DNA damage in LMNA-deficient hearts perturbs cell cycle and causes aberrant beating

Excess DNA damage has been shown to impact cell cycle in post-natal CMs (Puente, et al., 2014), and so we next sought to assess the biological consequences of DNA damage in LMNA-suppressed embryonic hearts. Morpholino-mediated knockdown of LMNA (‘MO_LMNA_’; ~40% KD in 24h) was achieved with no significant effect on contractile beating (**Fig.1F-*i***, **S1E**). LMNA is thus not primarily upstream of beating, consistent with knockout mice (Singh, et al., 2013). Although past studies also suggest LMNA is not detectable in early embryonic hearts and is therefore dispensable (Solovei, et al., 2013; Stewart and Burke, 1987), morpholino-knockdown increased γH2AX in E4 hearts (**Fig.1F-*ii***). Similar effects were seen by modest repression with retinoic acid (RA) of LMNA (−20% only after 3d) that acts transcriptionally (Swift, et al., 2013), while RA-antagonist (AGN) upregulated LMNA and suppressed DNA damage (**Fig.S1F,G**). Blebbistatin was sufficient to rescue the DNA damage in MO_LMNA_ knockdown hearts, supporting our hypothesis that high contractility coupled to low LMNA causes excess damage.

*Bona fide* DNA damage can perturb cell cycle (Puente, et al., 2014); the cell cycle inhibitor p21 indeed trended with γH2AX in MO-treated hearts (Fig.1F-*ii*). EdU integration before and after blebbistatin/washout (per Fig.1A-C) further revealed an increase in cells in G2, with fewer cells in S, and an increase in 4N cells (**Fig.1G**). Given that acute DNA damage can inhibit cell cycle in developing heart and is somehow coupled to matrix-myosin-lamins (**Fig.1H**), we assessed DNA damage that was directly and abruptly increased with etoposide (4-fold, 1h) (**Fig.2A**). This inhibitor of replication/transcription quickly caused aberrant beating that could be reversed upon drug washout (**Fig.2B**). No such effects were seen with the same dose of H2O2 (for oxidative stress) relative to DMSO control hearts, unless H2O2 exposure was sustained for days. DNA damage can have multiple causes, ranging from drugs and oxidative stress to nucleases, but reversal of damage from natural sources or etoposide in hours (Fig.1F,G,2A) suggest an important role for DNA repair factors (**Fig.2C**).

**Figure 2.**
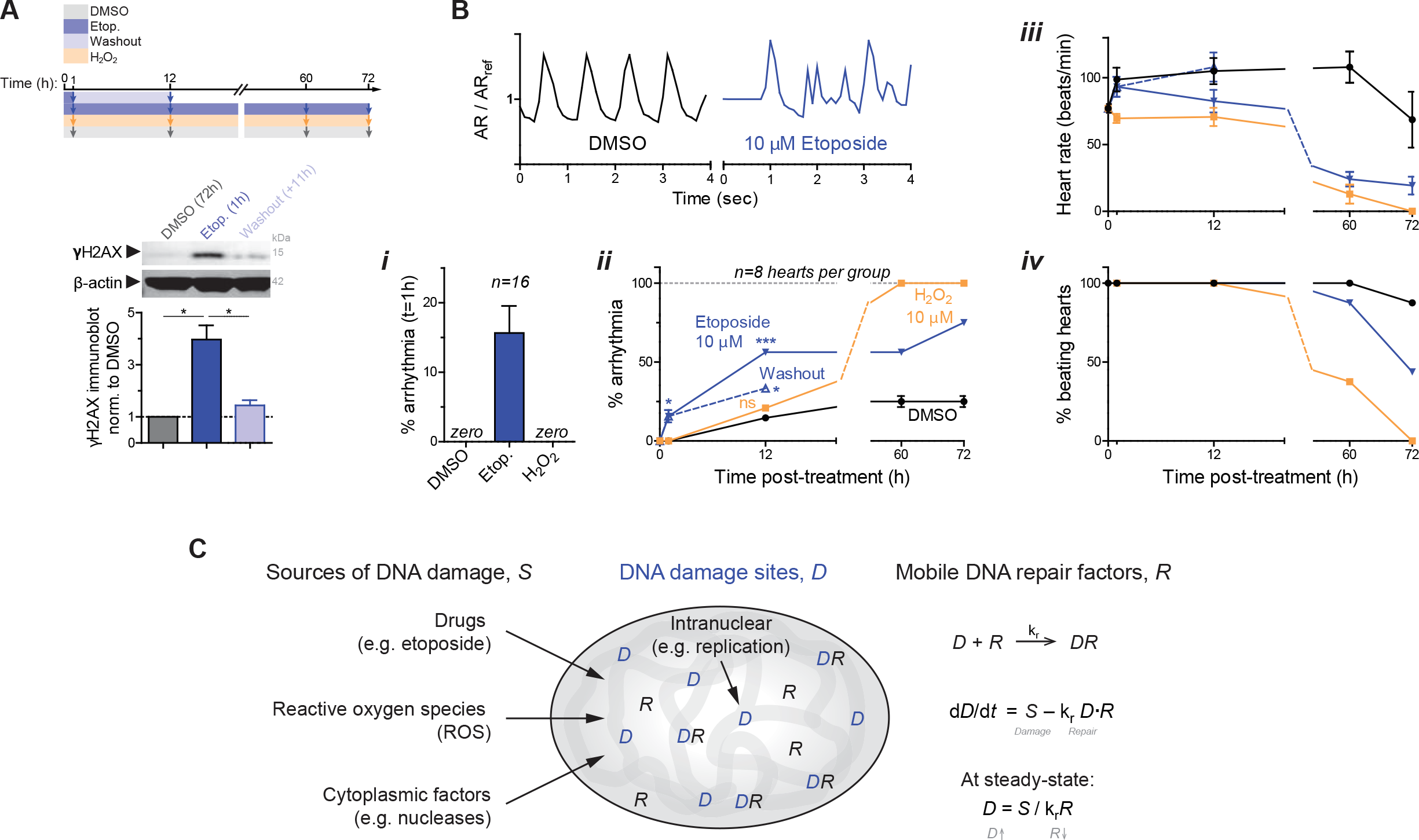
DNA damage in embryonic hearts causes aberrant beating. **(A)** 1h etoposide reversibly increases DNA damage 4-fold (8 hearts per grp). All error bars indicate ±SEM. **(B)** Beating (top) after 1h of treatment is only affected by etoposide. **(*i,ii*)** %-hearts exhibiting arrhythmia. Oxidant H2O2 shows distinct kinetics. **(*iii,iv*)** Beating decreases and arrests compared to DMSO. 8~16 hearts per grp; t-test \**p*<0.05, **<0.01, ***<0.001. **(C)** Schematic: the level of DNA damage that results from various sources depends also on levels of repair factors in the nucleus.

### LMNA knockdown in intact hearts & human iPS-CMs: nuclear rupture & loss of DNA repair

To address how increased mechanical stress causes DNA damage, the integrity of the nucleus was scrutinized in intact E5 hearts doubly transfected with DNA repair protein GFP-KU80 (XRCC5) and with a cytoplasmic protein that can bind nuclear DNA, mCherry-cGAS (cyclic GMP–AMP synthase (Raab, et al., 2016)) (**Fig.3A-*i***). Cytoplasmic mis-localization of KU80 *and* concomitant formation of cGAS puncta at the nuclear periphery provided evidence of nuclear rupture in transfected hearts (Fig.3A-*i*, yellow arrow), and the fraction of such double-positive cells increased with MO_LMNA_ knockdown unless blebbistatin was added (**Fig.3A-*ii***). Myosin-II inhibition disrupted sarcomeres, which rapidly recovered upon drug washout (**Fig.3A-*iii***), consistent with beating (Fig.1A-C); and mis-localization of KU80 and cGAS showed the same trends (**Fig.3A-*iv***), without any apparent changes in cell viability (**Fig.S2A-*i***). DNA damage again spiked upon blebbistatin washout and remained excessively high with addition of a nuclear import inhibitor (‘Imp-i’, ivermectin) that keeps KU80 mis-localized to cytoplasm (**Fig.3B-*i,ii***, **S2A-*ii***). LMNA immunofluorescence (IF) for MO_LMNA_ and blebbistatin/washout hearts (**Fig.3B-*iii* & C**) confirmed immunoblot trends (Fig.1C-*i*, F) and showed Imp-i kept LMNA levels low upon washout.

**Figure 3.**
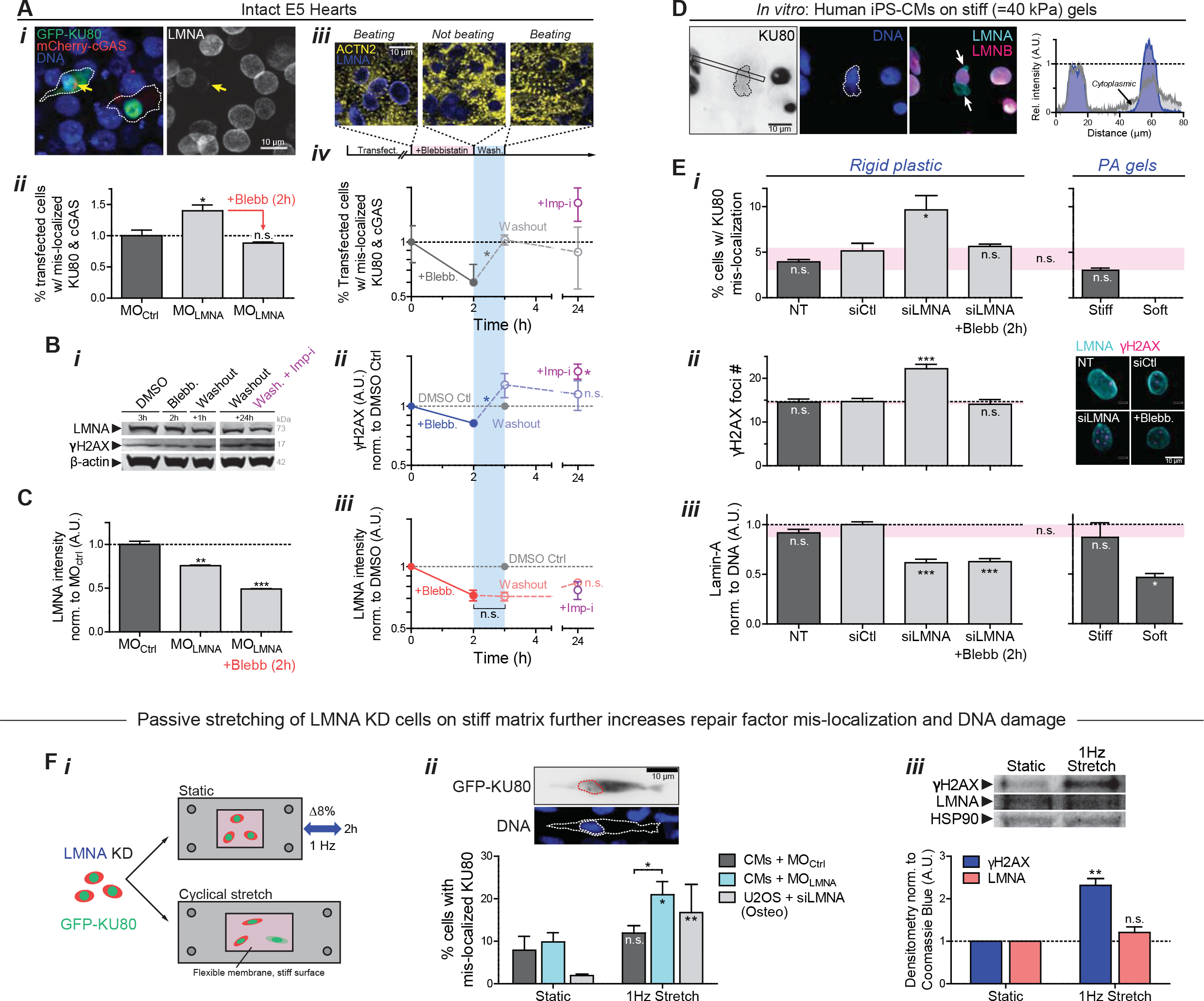
LMNA knockdown in intact hearts and in hiPS-CMs: nuclear envelope rupture and loss of DNA repair factors. **(A) (*i*)** Images of E4 hearts transfected with GFP-KU80 and mCherry-cGAS. Arrow: cell with low LMNA and ruptured nucleus, with cytoplasmic mis-localization of GFP-KU80 *and* mCherry-cGAS puncta at nuclear envelope. **(*ii*)** LMNA KD increases mis-localized KU80 and cGAS but is rescued by myosin-II inhibition. **(*iii*)** Images of striation (α-actinin-2) before blebb, 2h after, and upon washout (1h). **(*iv*)** %-Ruptured nuclei (double +’s) decreases with myosin-II inhibition but increases upon washout. Mis-localized KU80 increases further with import inhibitor ivermectin, Imp-i. (n>111 double-transfected cells per cond’n; t-test: \**p*<0.05). All error bars indicate ±SEM. **(B) (*i*,*ii*)** anti-γH2AX and **(*i*,*iii*)** anti-LMNA immunoblot of blebb-treated E4 hearts after washout ± Imp-i (8 hearts per lysate; t-test: \**p*<0.05; superfluous blot lanes have been removed and is indicated by a blank space). **(C)** LMNA immunofluorescence with MO_LMNA_ ± blebb. (n>139 cells per cond.). **(D)** Human iPS-derived cardiomyocytes (hiPS-CMs) on stiff 40 kPa gels for 24h, showing cytoplasmic KU80 in cells with nuclei having LMNA-rich, LMNB-depleted blebs at high curvature sites (arrows). **(E) (*i*,*ii*)** hiPS-CM on soft 0.3 kPa gels minimizes rupture. On rigid plastic, siLMNA KD (~40%, n>119 cells per cond. **(*iii*))** increases mis-localized KU80 (*i*) and DNA damage by γH2AX foci (right images). (n>80 cells per cond.; Dunnett’s test vs NT Ctrl) **(F) (*i*)** Passive cyclic stretch of LMNA-knockdown cells expressing GFP-KU80 (8% strain, 1Hz) increases **(ii)** %-rupture with cytoplasmic KU80 (upper inset) and **(*iii*)** γH2AX. (n>55 cells per cond.; t-test: #*p*<0.05, ##<0.01). (Unless noted,; one-way ANOVA: \**p*<0.05, **<0.01, ***<0.001)

Human induced pluripotent stem cell-derived CMs (hiPS-CMs) cultured on soft or stiff gels with collagen-ligand constitute another model for clarifying mechanisms at the single cell level. Nuclei in hiPS-CMs ‘beat’ (**Fig.S2B**) and express low LMNA relative to some well-studied human cell lines (e.g. lung-derived A549 cells) and relative to mature heart which is much stiffer than embryos (Swift, et al., 2013). Cell morphologies and sarcomere striations resemble those of embryonic/fetal CMs. On very stiff gels (40 kPa) but not on gels as soft as embryos (0.3 kPa), hiPS-CMs exhibited cytoplasmic, mis-localized KU80 (**Fig.3D**), consistent with envelope rupture under high nuclear stress (Fig.1H). Nuclear blebs typical of ruptured nuclei (Raab, et al., 2016) formed at points of high curvature in such cells (Xia, et al., 2018), appearing rich in LMNA but depleted of LMNBs (Fig.3D). Stress-induced nuclear rupture was also imaged as rapid and stable accumulation of mCherry-cGAS upon nuclear probing with a high curvature Atomic Force Microscopy (AFM) tip (<1μm), using nano-Newton forces similar to those exerted by a cell’s actomyosin (**Fig.S2C**).

KD of LMNA (‘siLMNA’) doubled the fraction of cells with cytoplasmic KU80 (**Fig.3E-*i***, **S2D-*i***) and increased γH2AX foci (**Fig.3E-*ii*** & **S2D-*ii***). LMNA levels in hiPS-CMs exhibited similar sensitivity to substrate stiffness as that in intact embryonic hearts (**Fig.3E-*iii***), and blebbistatin treatment in siLMNA cells on rigid plastic rescued nuclear rupture (Fig.3E-*i*) as well as DNA damage (Fig.3E-*ii*). Relaxation of rigidity-driven actomyosin stress can thus limit nuclear rupture.

Cytoplasmic mis-localization of KU80 is merely representative: repair factors 53BP1 and RPA2 also simultaneously mis-localized to the cytoplasm upon LMNA KD (**Fig.S2E-*i***). Time-lapse images of GFP-53BP1 expressing cells further revealed mis-localization persists for hours (**Fig.S2E-*ii***). However, total KU80 levels did not vary (**Fig.S2F**), suggesting nuclear retention is key.

To assess whether mis-localization of repair factors is a general consequence of increased mechanical stress, LMNA KD of both CM’s and U2OS osteosarcoma cells was followed by cyclic stretch at a frequency and magnitude similar to that of the intact heart (8% uniaxial strain at 1 Hz frequency) (**Fig.3F-*i***). Mis-localized GFP-KU80 increased >2-fold with imposed stretching of MO_LMNA_ knockdown E4 CMs but not MO_Ctrl_ (**Fig.3F-*ii***), and the fold-increase was even more dramatic with the non-beating U2OS cells because baseline mis-localization was very low. DNA damage also increased with stretching of MO_LMNA_ CMs while LMNA levels showed an insignificant increase (**Fig.3F-*iii***). An acute increase in stress without a proportionate increase in LMNA can thus cause mis-localization of repair factors and DNA damage, which could affect tissue-level function (Fig.2A,B). Etoposide treated hiPS-CM organoids indeed show aberrant beating (**Fig.S2G**).

### Loss of Repair Factors or LMNA: DNA damage, binucleation & micronuclei, cell cycle arrest

To assess the impact of a limited capacity for DNA repair in cells, key individual DNA repair factors were partially depleted individually and in combination (‘si-Combo’). BRCA1 is implicated in myocardial infarction and ischemia (Shukla, et al., 2011), and RPA1 appeared abundant and constant in level in our MS of drug/enzyme-perturbed hearts (Fig.1C,E). DNA damage increased in hiPS-CMs for all repair factor KDs, and the combination proved additive (**Fig.4A-*i***).

**Figure 4.**
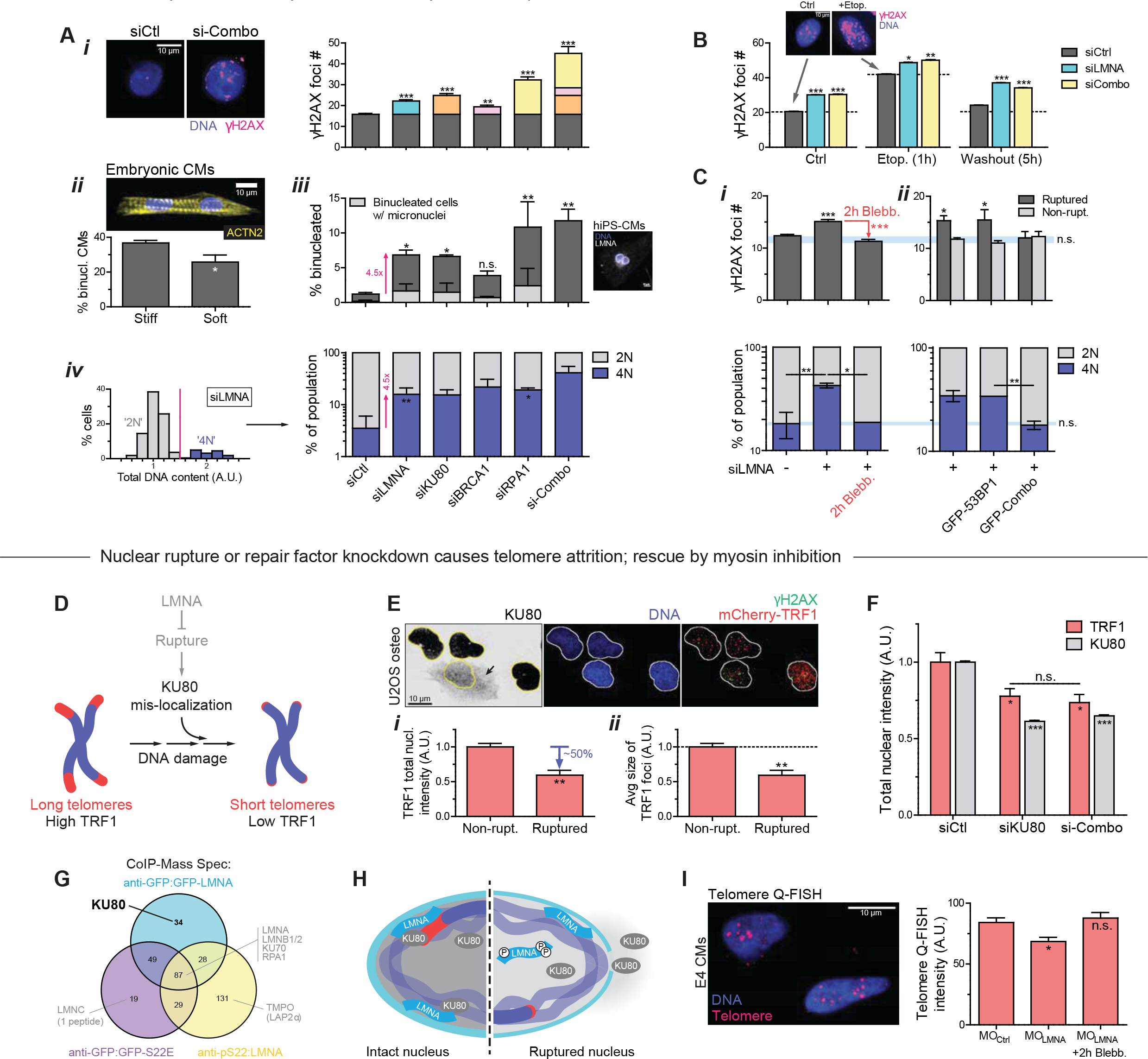
Knockdown of LMNA or DNA repair factors increases DNA damage and perturbs cell cycle. **(A)** KD of LMNA or repair factors (KU80, BRCA1, RPA1, or ‘si-Combo’) in hiPS-CMs increases: **(*i*)** γH2AX **(*ii*,*iii*)** binucleated CMs (suppressed on soft matrix, inset) and micronuclei, and **(*iv*)** 4N cells (>77 cells per cond.; all error bars indicate ±SEM) **(B)** KDs increase γH2AX before/during/after 10 μM etoposide (n>39 cells per cond). **(C) (*i*)** Blebb rescues siLMNA-induced DNA damage and 4N cells (n>400 cells per cond.). **(*ii*)** Over-expression of DNA repair factors (GFP-KU70 + KU80 + BRCA1 = ‘GFP-Combo’) rescues excess DNA damage (top) and 4N (bottom) in siLMNA cells only in ruptured nuclei (n>296 cells per cond.; t-test: \*\**p*<0.01, ***<0.001). **(D)** Schematic: effect of repair factor mis-localization on telomeres. **(E)** U2OS expressing mCherry-TRF1. Ruptured nuclei (n=7) with cytoplasmic KU80 exhibit **(*i*)** lower TRF1 and **(*ii*)** smaller TRF1 foci (t-test: *p<0.05). **(F)** siKU80 and si-Combo decrease TRF1 (n>89 cells per cond.). **(G)** CoIP-MS of WT LMNA, phosphorylated LMNA (anti-S22), and phospho-mimetic LMNA (GFP-S22E). **(H)** Cartoon: LMNA-KU80-telomere interactions at the lamina. **(I)** Telomere Q-FISH in embryonic CMs shows MO_LMNA_ shortens telomere unless rescued by blebb (n>81 nuclei per cond.). (Unless noted, one-way ANOVA: *p<0.05, ***<0.001)

Excess DNA damage in post-natal hearts has been associated with CM binucleation (Aix, et al., 2016) – a hallmark of irreversible cell cycle exit. Soft ECM suppressed binucleation of embryonic CMs relative to stiff ECM (**Fig.4A-*ii***) while suppressing mis-localization of repair factors (Fig.3E-*i*). Partial KD of DNA repair factors directly increased the fraction of binucleated hiPS-CMs as well as those with micronuclei (**Fig.4A-*iii***), both of which correlated strongly with γH2AX. Repair factor KD further increased the fraction of ‘4N’ cells (vs ‘2N’; **Fig.4A-*iv***, **S3A**), in agreement with trends for blebbistatin-treated hearts (Fig.1G) and with mitotic arrest. LMNA KD had all of the same effects, consistent with a role in repair factor retention in nuclei, and consistent also with a late checkpoint, KD showed more DNA damage per DNA in ‘4N’ cells relative to siCtrl cells (**Fig.S3B**). Importantly, with siLMNA or siCombo KD, increased γH2AX and cell cycle perturbations were evident not only at steady state but also within 1hr of damaging DNA with etoposide (per Fig.2) – which partially recovered after drug washout (**Fig.4B**).

Myosin-II inhibition once again rescued the DNA damage as well as cell cycle effects in siLMNA KD cells (**Fig.4C-*i***), whereas siCombo cells maintained high DNA damage (**Fig.S3C**). Furthermore, over-expression of relevant DNA repair factors (with GFP-KU70, KU80, BRCA1 = ‘GFP-Combo’) rescued only the excess DNA damage in nuclear-ruptured cells compared to non-ruptured (**Fig.4C-*ii***). A baseline level of damage in the latter cells was not affected, likely because repair factors become limiting only upon nuclear rupture or depletion. A combination of factors (‘GFP-Combo’) rescued the excess damage while one repair factor had no effect on its own (53BP1) even on the ruptured nuclei. Rescue of DNA damage again suppressed the fraction of ‘4N’ cells (Fig.4B lower) and cells in G2 (**Fig.S3D**).

Since cardiac telomere attrition has been implicated in CM binucleation (Aix, et al., 2016) as well as in cardiomyopathies and heart failure (Sharifi-Sanjani, et al., 2017), we assessed rupture-associated telomere shortening as a more localized form of DNA damage (**Fig.4D**). U2OS osteosarcoma cells with ruptured nuclei (cytoplasmic anti-KU80) showed lower total intensity of TRF1 (telomeric repeat-binding factor-1 as an mCherry fusion; **Fig.4E-*i***) and smaller TRF1 foci size (**Fig.4E-*ii***), as well as higher γH2AX in these LMNA KD cells. Importantly, partial KD of either KU80 or multiple repair factors also caused telomere attrition in cells with normal LMNA levels (**Fig.4F**). Proper retention of DNA repair factors within stressed nuclei thus requires sufficient LMNA, but LMNA also interacts with key repair factors (including KU70/80, RPA1) as confirmed by immunoprecipitation-MS (IP-MS) (**Fig. S3E**). IP-MS further indicates KU80 interacts preferentially with non-phosphorylated LMNA (’GFP-WT’) compared to its phospho-solubilized nucleoplasmic form (per IP by anti-pSer22 or phosphomimetic mutant ‘GFP-S22E’) (**Fig.4G**), suggesting that LMNA scaffolds the repair of DNA at the periphery – which often includes telomeres (**Fig.4H**). Importantly, analysis of telomeres in embryonic CMs by quantitative FISH (Q-FISH) revealed LMNA KD causes telomere shortening, unless actomyosin is inhibited (**Fig.4I**). The findings motivate deeper mechanistic insights into how LMNA adjusts to actomyosin stress (Fig.1A-E & 2C, 2E-*iii*) to thereby regulate DNA damage and cell cycle.

### LMNA effectively stiffens nuclei during development and increases with collagen-I ECM

To begin to assess in normal beating hearts how LMNA senses stress, nuclear deformations were imaged at embryonic day-4 (E4) – just 1~2d after the first heartbeat (**Fig.5A**). Transfection of a minor fraction of CMs in hearts with mCherry-Histone-H2B or GFP-LMNA facilitated imaging, and each ~1Hz contraction of heart tissue was seen to strain individual CM nuclei by up to ~5-8%. Recall that 8% stretch at 1-Hz was sufficient to rupture CM nuclei (Fig.3F). While transfection did not affect overall tissue-level beating, GFP-LMNA nuclei deformed much less than neighboring control nuclei (**Fig.5A-*ii***), consistent with *in vitro* studies showing LMNA stiffens the nucleus (Pajerowski, et al., 2007; Lammerding, et al., 2006). MS of lysates quantified 29 unique LMNA peptides at E4 (Fig.S1C), confirming our early detection of functional LMNA (Fig.1B) despite past reports (Solovei, et al., 2013; Stewart and Burke, 1987), and showed a rapid increase in LMNA expression by ~3-fold from soft E4 hearts to stiffer E10 hearts (**Fig.5B,C**). B-type lamins remained relatively constant, in agreement with studies of diverse soft and stiff adult tissues that suggest LMNA-specific mechanosensitivity (**Fig.S4A-C**) (Swift, et al., 2013). Trends are confirmed by confocal IF and immunoblots of embryonic heart (**Fig.5C-*i*,*ii*** **insets**, **S4D**).

**Figure 5.**
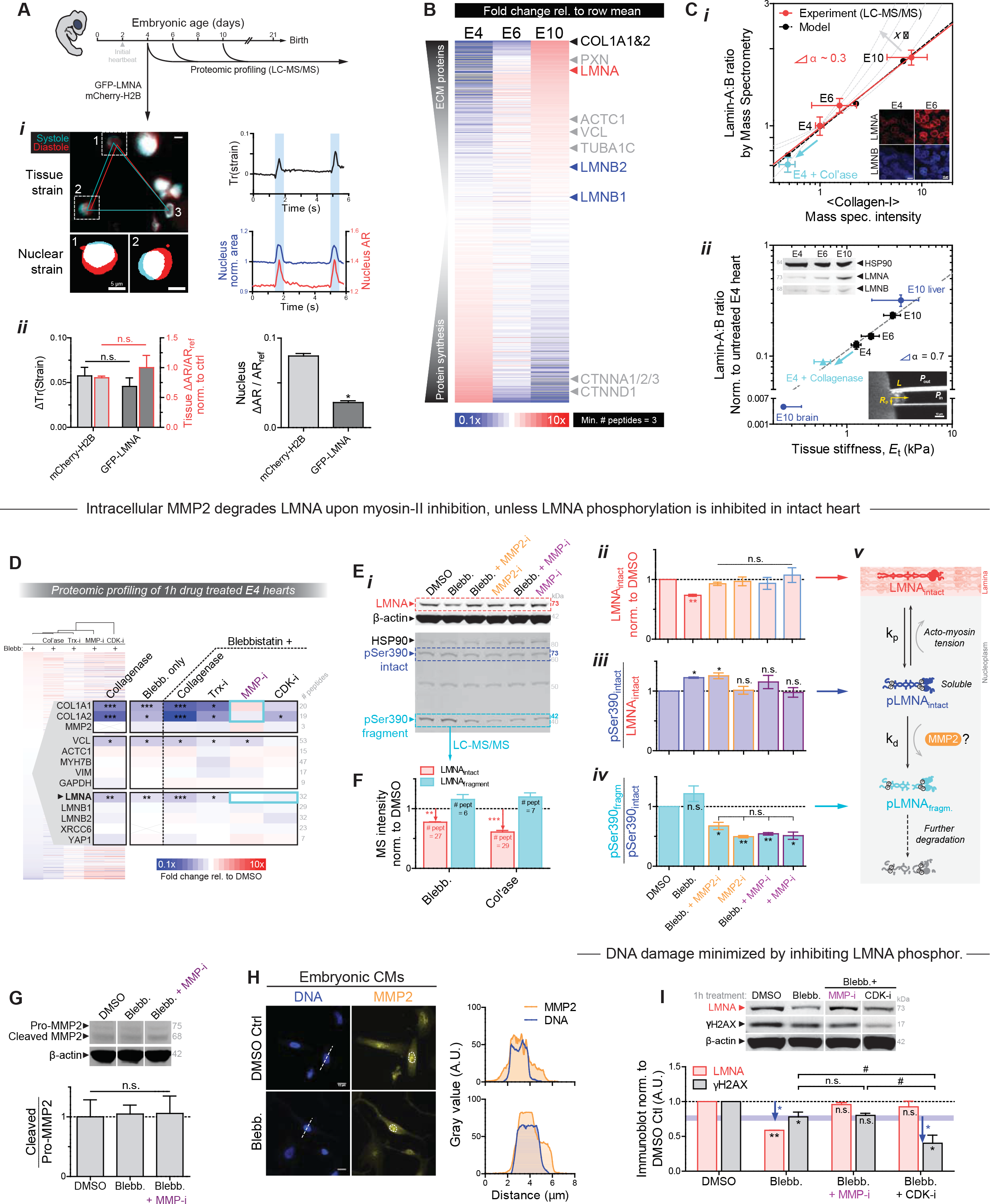
Early embryonic LMNA increases with collagen-I and tissue stiffness *E*_t_. **(A) (*i*)** E4 heart beating with GFP-LMNA transfection shows nuclear ‘beating’. Right: Trace of tissue strain tensor quantified using fiduciary transfected nuclei per (Taber, et al., 1994), or nuclear strain from nuclear area and aspect ratio (AR). **(*ii*)** Overexpression does not affect local or global tissue strain (left), but GFP-LMNA nuclei deform less than mCherry-H2B nuclei (n>8 nuclei per cond.; t-test: \**p*<0.05). All error bars indicate ±SEM. **(B)** Proteomic heatmap of E4, 6, and 10 hearts (n=8,2,1 hearts per lysate, resp.), ranked by fold-change from time-average: collagen-I and LMNA top the list. **(C) (*i*)** Lamin-A:B ratio from MS scales with collagen-I and fits ‘lose it or use it’ model of tension-inhibited turnover of structural proteins, with stress-sensitivity denoted as x. Inset: Confocal images of LMNA & B in hearts taken at same settings suggest LMNA increases while LMNB remains constant. **(*ii*)** Lamin-A:B scaling vs *E*_t_ from micropipette aspiration (lower inset), with exponent α similar for adult tissue (α ~ 0.6) (Swift, et al., 2013). Cyan: softening by collagenase decreases LMNA. Upper immunoblot (norm. to HSP90): LMNB varies by <20% from E4-E10 as LMNA increases (n>8 hearts per lysate). **(D)** MS intensity fold-changes relative to DMSO for ‘Trx-i’ inhibitor of translation (cycloheximide, 1μM); ‘MMP-i’ pan-MMP inhibitor (GM6001, 10μM); ‘CDK-i’ (RO3306, at >3.5 μM doses to inhibit many CDKs) to suppress LMNA phosphorylation. Cyan boxes indicate rescues (n=8 hearts per lysate; t-test vs DMSO: \**p*<0.05, **<0.01, ***<0.001). **(E) (*i*)** E4 hearts treated with blebb ± MMP-i’s (2h; n=6 hearts per lysate). pSer390 bands at intact MW (73 kDa) and 42 kDa, consistent with fragments (Baghirova, et al., 2016; Prudova, et al., 2010) predicted *in silico* (Song, et al., 2012). **(*ii*)** Decreased LMNA with blebb is rescued by MMP-i’s, including MMP2-specific ARP-100. **(*iii*)** Blebb increases pSer390 of 73 kDa band. **(*iv*)** Blebb does not affect pSer390 ratio of 42 kDa and 73 kDa bands, except with MMP-i. (n=7 hearts per lysate; one-way ANOVA: \**p*<0.05, ** *p*<0.01) **(*v*)** Turnover schematic. **(F)** MS validation of LMNA degradation. **(G)** Neither blebb nor MMP-i affect catalytic activation of MMP2 (n=8 hearts per lysate). **(H)** IF shows nucleoplasmic MMP2 in embryonic CMs is unaffected by blebb. **(I)** MMP-i rescues blebb-induced decrease in LMNA, but does not further suppress γH2AX (n=6 hearts per lysate). γH2AX is further decreased with CDK-i to inhibit LMNA phosphorylation (n=8 hearts per lysate; one-way ANOVA: \**p*<0.05, **<0.01; t-test: #p<0.05).

Collagen-I (α1, α2) exhibited the largest fold-change in MS analyses of heart development (**Fig.5B**), with calibrated spike-ins of purified collagen-I matching past measurements of collagen in developing chick hearts (Woessner, et al., 1967) (**Fig.S4E**). Collagen-I becomes the most abundant protein in developed animals and is a major determinant of tissue stiffness (Shoulders and Raines, 2009), and so its increase together with many other structural proteins (**Fig.S4C**) is consistent with heart stiffening (**Fig.S2F**) (Majkut, et al., 2013). Mechanosensitive proteins at the interface between adhesions and actomyosin cytoskeleton were also upregulated, including paxillin (Zaidel-Bar, et al., 2007) and vinculin (Huang, et al., 2017), consistent with increased adhesion and contractile stress (Geiger, et al., 2009). Down-regulation of protein synthesis and catenins suggest a transition from a rapidly growing, epithelial-like embryo to a more specialized, terminally differentiated state.

Lamin-A:B ratio increased with collagen-I (**Fig.5C-*i***, S4F) and with tissue stiffness *E*_t_ (**Fig.5-*ii***), with a power law exponent α=0.3 (vs collagen-I) similar to that for diverse tissue proteomes (Cho, et al., 2017). Experiments fit a ‘lose it or use it’ mathematical model (Fig.5C-*i*) wherein mechanical tension on filamentous collagen-I and LMNA (Dingal and Discher, 2014) suppresses protease-driven degradation (Fig.5C-*i*). The plot versus *E*_t_ yielded α~0.7 that is similar for adult tissues (α~0.6 (Swift, et al., 2013)), with soft E10 brain and stiff E10 liver fitting well (**Fig.S4F,G**). Importantly, 45min collagenase softened E4 hearts by ~50% and decreased LMNA (Fig.5C), per Fig.1A-E.

Different combinations of drugs were used to identify mechanisms for how LMNA levels change in tight coordination with collagen-I and tissue stiffness. MS-derived heatmaps of drug-treated E4 hearts (e.g. collagenase, blebb) reveal rapid and large decreases in collagen-I, vinculin, and LMNA (**Fig.5D**; confirmed by immunoblot, **Fig.S5A**). Little to no change was observed across most of the detected proteome including DNA repair factors (KU70, RPA1 (**Fig.S5B-*i***)), matrix-metalloproteinase MMP2 (pro- and cleaved forms, **Fig.S5B-*ii*,*iii***), cardiac myosin-II (MYH7B), vimentin (VIM), and B-type lamins. Similar results were obtained even in the presence of the translation inhibitor cycloheximide (‘Trx-i’), except for decreases in a few cytoskeletal proteins that indicate rapid transcription/translation (e.g. actin (Katz, et al., 2012)). Decreased translation thus does not explain the heart’s decreases in collagen-I and LMNA with blebbistatin and/or collagenase.

To clarify how myosin-II inhibition (intracellular) leads to a rapid decrease in collagen-I (ECM) in intact hearts, a pan-inhibitor of matrix-metalloproteinases (‘MMP-i’) was used with blebbistatin. The combination surprisingly prevented collagen-I degradation. In contrast, a CDK inhibitor (‘CDK-i’, likely affecting CDK4/6) that limits interphase phosphorylation of LMNA (e.g. pSer22 normalized to total LMNA: ‘pSer22/LMNA’ in Fig.S5A) rescued blebbistatin-induced decreases in LMNA levels without affecting the collagen-I decrease. Vinculin’s decrease concurs with decreased actomyosin tension regardless of MMP-i (Fig.5D), but MMP-i surprisingly rescued blebbistatin-induced decreases in LMNA similar to CDK-i (Fig.5D, S5A).

### Mechano-protection of the genome is limited by LMNA phospho-solubilization

We hypothesized that one or more MMPs directly degrade LMNA. Nuclear MMP2 is well-documented for many cell types including CMs (Xie, et al., 2017), and LMNA (but not B-type lamins) has been speculated to be a proteolytic target (Baghirova, et al., 2016). Beating E4 hearts treated with MMP-i or an MMP2-specific inhibitor, MMP2-i, rescued the blebbistatin-induced decrease in LMNA in immunoblots that were also probed with a novel anti-pSer390 (**Fig.5E-*i,ii***). For other phosphosites, interphase phosphorylation, solubilization (Kochin, et al., 2014), and subsequent degradation of LMNA (Naeem, et al., 2015; Bertacchini, et al., 2013) are already known to be suppressed by actomyosin tension on nuclei (Buxboim, et al., 2014; Swift, et al., 2013). Intact phospho-LMNA (73 kDa) increased upon blebbistatin treatment (**Fig.5E-*iii***, ‘pSer390_intact_/LMNA_intact_’), but even more dramatic was the suppression by MMP2-i and MMP-I (regardless of blebbistatin) of a 42 kDa pSer390 fragment (‘pSer390_fragm._/pSer390_intact_’) (**Fig.5E-*iv***). Fragment size matches predictions for MMP2 digestion (Song, et al., 2012), but is likely a transient intermediate that is further degraded (Buxboim, et al., 2014) (**Fig.5E-*v***). MS of hearts treated with blebbistatin or collagenase not only confirmed the trend for the fragment (>6 LMNA peptides) (**Fig.5F**) but also showed MMP2 levels remain constant, as supported by immunoblots for MMP2’s activation (‘cleaved/pro-MMP2’ ratio, **Fig.5G**, S5B). Anti-MMP2 (for both pro- and active forms) was mostly nucleoplasmic in isolated CMs with localization unaffected by actomyosin inhibition (**Fig.5H**). Nucleoplasmic MMP2 in neonatal CMs (Baghirova, et al., 2016; Kwan, et al., 2004) suggests that, independent of maturation stage, active MMP2 is localized appropriately for degradation of phospho-LMNA downstream of actomyosin tension (Fig.5E-*v*).

DNA damage was assessed after acute 2h treatments of CDK-i/MMP-i plus blebbistatin. Whereas MMP-i had no effect on γH2AX relative to blebbistatin alone, CDK-i plus blebbistatin greatly suppressed DNA damage (**Fig.5I**). Since CDK-i maintains LMNA at the lamina (unlike MMP-i) and thereby stiffens and stabilizes the nucleus (per Fig.5A and single cell measurements (Buxboim, et al., 2014)), the results suggest that even low levels of nuclear stress are resisted to minimize both rupture and DNA damage in the embryonic heart.

### Isolated CMs establish ECM elasticity and contractility in LMNA turnover by MMP2

Intact heart presents many complications to understanding LMNA regulation and its function. As the heart stiffens in development, for example, collagen ligand also increases greatly for adhesion (Fig.5B,C), sarcomere assembly becomes very dense (**Fig.6A**, **S6A-*i***), cells and nuclei elongate (**Fig.S6A-*ii***, **S6B**) with stabilization by microtubules (Robison, et al., 2016), cells align (**Fig.S6C**), and nuclear volumes decrease (**Fig.S6D**). Gels with constant collagen-I ligand and of controlled stiffness (0.3 - 40 kPa) (**Fig.6B**) showed – after just 24h – the same stiffness-correlated morphology and cytoskeleton trends for isolated E4 CMs as the intact heart from E4 to ~E11 (**Fig.6C**). Although matrix elasticity had large effects on size, shape, sarcomere assembly, and contractility of early CMs (Fig.S6A-*i*,*ii*, **S6E**) similar to later stage CMs (Ribeiro, et al., 2015; Engler, et al., 2008; Jacot, et al., 2008), blebbistatin quickly disrupted cell spreading, striation, and elongation on stiff gels (10 kPa) and on collagen-coated rigid plastic (**Fig.6C**). Gels that mimic the stiffness of E4 heart (1-2 kPa) were optimal for cell beating (**Fig.S6F**), consistent with past studies (Majkut, et al., 2013; Engler, et al., 2008), and nuclear ‘beating strain’ in isolated CMs (Fig.S6F-*i*, **S6G-*i***) was similar in magnitude to that in intact heart (5-8%) (Fig.5A). Surprisingly, lamin-A:B intensity increased monotonically with gel stiffness rather than exhibiting an optimum (**Fig.6D-*i***, **S6G-*ii***). LMNA’s increase depended on myosin-II, as did cell/nuclear aspect ratio (AR), spreading, and sarcomere assembly (Fig.6C, **5D-*ii***, **S6H**). Monotonic increases with stiffness of ECM agrees with elevated isometric tension as documented for other cell types (e.g. (Engler, et al., 2006)). Remarkably, MMP-i or MMP2-i rescued the blebbistatin-induced decrease in lamin-A:B at the single cell level (Fig.5D-i) as seen for intact embryonic hearts (Fig.5D-i), despite no significant effects on cell morphology or striation (Fig.6C,D-*ii*).

**Figure 6.**
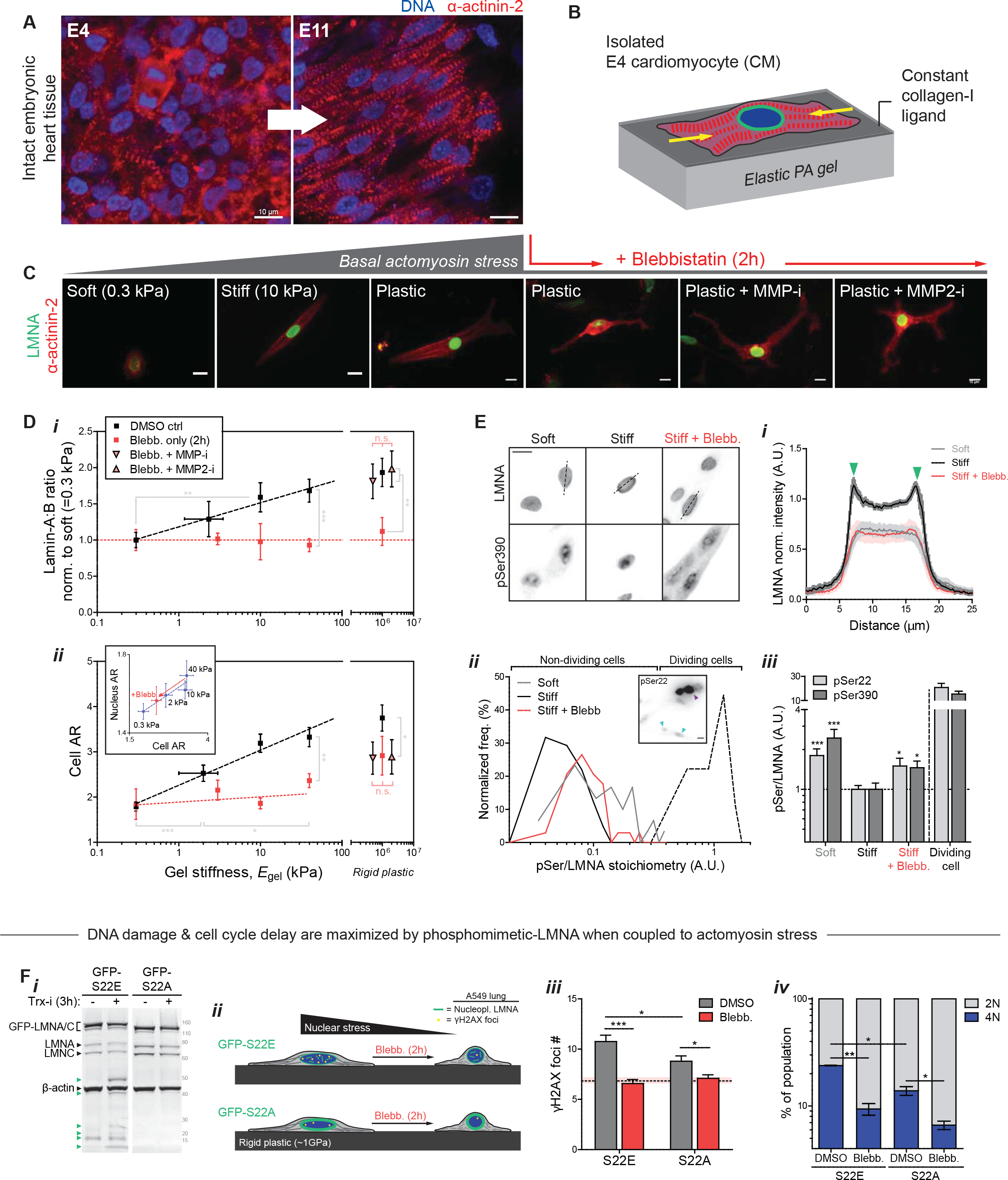
LMNA in isolated embryonic CMs is sensitive to substrate stiffness and actomyosin stress. **(A)** Airyscan images of embryonic heart. **(B)** CM cultures on PA gels of controlled stiffness (0.3 −40 kPa) and constant collagen-I ligand. **(C)** Well-separated E4 CMs at 24h on gels. As with tissue (A), isolated CMs spread, elongate, and striate more with stiffness. Blebb on rigid plastic causes rounded or dendritic CM, with loss of striation. MMP inhibitors have no effect. **(D) (*i*)** Lamin-A:B increases with gel stiffness except with blebb (n>50 cells per grp). Effects are rescued by MMP-i’s on rigid plastic. **(*ii*)** Cell and nuclear AR (inset) increase as CMs polarize on stiffer gels. Blebb reduces cell/nuclear AR, but MMP-i’s show not affect (n>47 cells per cond.; all error bars indicate ±SEM). **(E) (*i*)** LMNA is more nucleoplasmic in CMs on soft gels (0.3 kPa) than stiff (40 kPa). Blebb gives higher nucleoplasmic signal. **(*ii*)** Histograms of phosphorylation (‘pSer/LMNA’) for pSer22 and pSer390. Signal for both sites are higher for soft gels and stiff gels + blebb. pSer/LMNA for rare mitotic CMs (yellow) are typically 10~20-fold higher than interphase CMs (cyan arrows). **(*iii*)** pSer/LMNA ratios for pSer22 and 390 (n>11 nuclei per cond.). **(F) (*i*)** Phospho-mimetic mutant GFP-S22E (± Trx-i, 3h) probed with anti-LMNA shows low LMNA versus GFP-S22A cells. Stable lines were made within a stable shLMNA line to minimize both endogenous LMNA and overexpression artifacts. Low-MW fragments (green arrows) are not detected in GFP-S22A cells. **(*ii,iii*)** Basal DNA damage and **(*iv*)** %4N cells are highest in GFP-S22E cells, but blebb minimizes differences (n>93 nuclei per cond.; t-test \**p*<0.05, **<0.01, ***<0.001; irrelevant blot lanes have been removed and is indicated by a blank space).

Since myosin-II-dependent increases in lamin-A:B versus gel stiffness for isolated CMs (Fig.6D-*i*) align with lamin-A:B’s increase with heart stiffness and contractility during development (Fig.5B,C), LMNA phosphorylation was re-examined (per Fig.5D, S5A). IF with anti-pSer22 & pSer390 proved consistent with tension-suppressed LMNA phosphorylation and turnover: cells on soft gels showed high nucleoplasmic phospho-signal as did cells on stiff gels treated with blebbistatin (**Fig.6E**). The latter also showed cytoplasmic phospho-signal, consistent with multi-site phosphorylation in interphase (Kochin, et al., 2014). Regardless, LMNA was similarly low and decreased at the nuclear envelope (**Fig.6E-*i***). Phospho-signal normalized to total LMNA (‘pSer/LMNA’) was also 2~3-fold higher in CMs on soft gels compared to stiff gels (**Fig.6E-*ii,iii***). Blebbistatin-treated cells on stiff gels again showed increased pSer/LMNA for both phospho-sites. Signal in interphase cells was, however, >10-fold lower than in rare mitotic CMs (Fig.6E-*ii*, inset: purple arrow). Studies of stable phospho-mimetic mutants (GFP-S22A & GFP-S22E) conformed to the model, with higher nucleoplasmic signal but lower overall levels for S22E (**Fig.6F-*i***, **S7A**). Inhibition of translation for 3 hrs (Trx-i) showed more degradation only for S22E (Fig.6F-*i* **S7B**), and MMP2-i treatment increased S22E intensity (**Fig.S7C**) consistent with inhibition of phospho-LMNA turnover. S22E-phosphomimetic cells also exhibited higher DNA damage (**Fig.6F-*ii,iii***) and more cells in late cell cycle (‘4N’, **Fig.6F-*iv***) that were more suppressed by acute blebbistatin than S22A cells. Reduction of nuclear stress by soft matrix or actomyosin inhibition at the single cell level thus favors LMNA interphase phosphorylation, nucleoplasmic solubilization, and subsequent degradation – while also suppressing DNA damage and cell cycle arrest.

## Discussion

LMNA defects cause disease through “cell-extrinsic mechanisms” that likely include ECM and/or cytoskeletal stress. Mosaic mice in which 50% of the cells express defective LMNA maintain a normal lifespan, whereas mice with 100% defective cells die within weeks of birth (de La Rosa, et al., 2013). Cultures on rigid plastic of the same cells (and similar cell types (Hernandez, et al., 2010)) exhibit premature senescence/apoptosis, as is common with excess DNA damage, but growth and viability are surprisingly rescued upon culture on almost any type of ECM. Soft matrix certainly reduces cytoskeletal stress and suppresses nuclear rupture and DNA damage in CMs with low LMNA (Fig.3E). The findings are further consistent with the observation that laminopathies spare soft tissues such as brain, independent of lineage or developmental origin, but generally affect stiff and mechanically stressed adult tissues including muscle or bone (Cho, et al., 2018).

Early embryos are soft, with minimal ECM, but as tissues form and sustain higher physical stresses, collagen-I accumulates and stiffening of the developing tissue minimizes the strain on resident cells. However, increasing stiffness and stress necessitate an adaptive mechanism to protect against nuclear stress, and accumulation of mechanosensitive LMNA fulfills this role – unless actomyosin stress is inhibited (Fig.1A-C). LMNA thus mechano-protects DNA by retaining repair factors in the nucleus (Fig.3A-D), which thereby prevents excessive DNA damage in stiff microenvironments and/or under high actomyosin stress (**Fig.7A**). Rapid change in LMNA protein independent of any transcription/translation (Trx-i) rules out many possible contributing pathways to the equally rapid DNA damage response. Such conclusions about causality are difficult to otherwise achieve with experiments conducted over many hrs/days such as with mouse models. Results with the cardiac myosin-II specific inhibitor (‘MYK’) in clinical trials for some cardiomyopathies (NCT02842242) further suggest a novel means to attenuate DNA damage in heart (Fig.1C,D).

**Figure 7.**
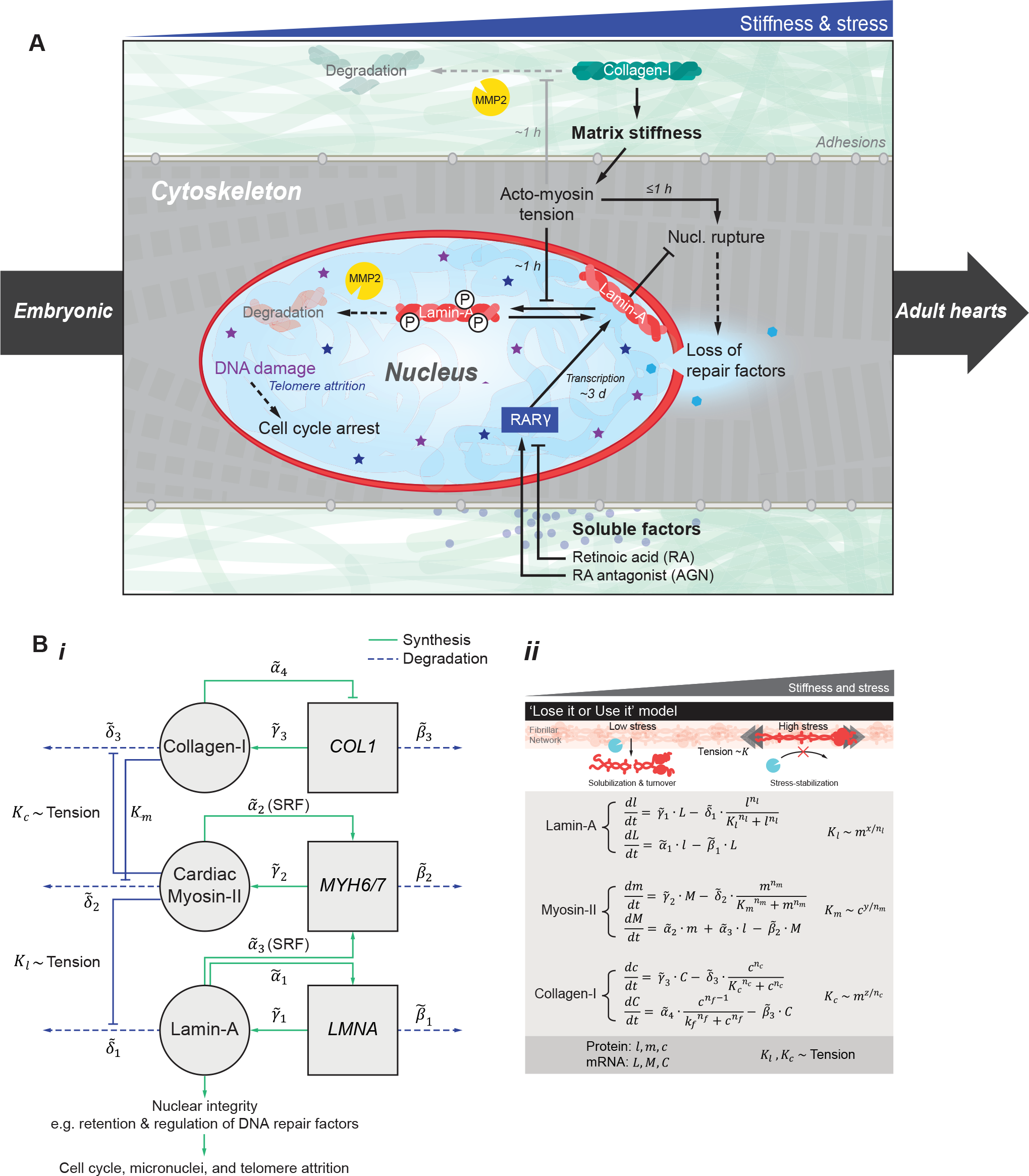
LMNA mechanosensing based on a ‘lose it or use it’ mechanism for tension-inhibited turnover that also applies to collagen-I. (A) Summary sketch. (B) **(*i*)** Gene circuit model adapted from Dingal et al. (Dingal and Discher, 2014), with (*ii*)governing equations. Squares = genes, circles = protein. LMNA and myosin-II protein (*l* and *m*) are weak regulators of the Serum Response Factor (SRF) pathway. LMNA upregulates its own transcription (via RARγ). LMNA and collagen-I protein turnover with *x* and *z* sensitivity of degradation to myosin tension. Myosin-II protein turnover depends on matrix elasticity *E* which scales with collagen-I (Fig.S4E).

Loss of DNA repair factors could compromise genome integrity, but enhanced entry of cytoplasmic nucleases (Maciejowski, et al., 2015) would seem unlikely to generate γH2AX foci distributed throughout the nucleoplasm (Fig.3E-ii, 4D). Indeed, cytoplasmic proteins that bind DNA strongly (e.g. cGAS) are restricted to rupture sites. Rupture results in ‘global’ mis-localization of multiple DNA repair factors in several pathways (Fig.S2E-i) (Xia, et al., 2018; Irianto, et al., 2017), and repair factor depletion causes excess DNA damage before, during, and after transiently induced damage (by 1h etoposide, Fig.4B).

Remarkably, MMPs that degrade and remodel collagen-I matrix within hours or less are found here to directly regulate LMNA of the ‘nuclear matrix’ (Fig.5D-I, 5A,B), which indicates a surprising inside-outside symmetry (Fig.7) to the ‘lose it or use it’ mechanism (Fig.5C, **7B**) of stress-stabilizing fibers. While both phospho-LMNA and MMP2 are found mostly in the nucleoplasm of CMs, the specific functions of MMP2 in the nucleus, the location of LMNA degradation, and its activation/localization mechanisms remain to be elucidated. Nuclear MMPs are not new (Xie, et al., 2017), but tension-regulation of their substrates has not been described. Our studies of isolated CMs on soft/stiff gels further demonstrate steady-state LMNA levels are determined primarily by a basal tone related to average morphologies of cells and nuclei, as opposed to dynamic contractions that intermittently strain the nucleus (Fig.3F, 5D, S6G). Basal and dynamic strain might contribute to a ‘tension-time integral’ model (based on cumulative time spent under high tension) that also predicts hypertrophic versus dilated cardiomyopathy disease fates (Davis, et al., 2016). Roles for MMPs in such diseases and compensatory changes in LMNA could emerge.

Accumulation of LMNA in stiff tissues of the embryo occurs far earlier than previously reported (Stewart and Burke, 1987) and tracks non-linearly what eventually becomes the most abundant protein in adult animals, collagen-I (Fig.5C). Collagenase softens tissue and decreases LMNA consistent with scaling trends, and the scaling LMNA ~ collagen-I^0.3^ in embryos matches meta-analyses of all available transcriptomes analyzed for diverse normal and diseased adult and developing hearts across species (Cho, et al., 2017) (**Fig.S7A**). The scaling is consistent with that across proteomes of diverse chick embryo tissues at E18 (**Fig.S7B**) as well as adult mouse tissues (α=0.4) (Swift, et al., 2013), and underscores a universality of normal regulation. Even the tissue-dependent timing of LMNA detection (Solovei, et al., 2013; Rober, et al., 1989; Lehner, et al., 1987) correlates well with stiffness of a tissue in adults (**Fig.S7C**; adapted from (Swift, et al., 2013; Rober, et al., 1989)). Mechano-coupled accumulation of LMNA and collagens that begin in early development could thus persist well into maturation and perhaps even to disease and aging of adult tissues (Fig.7G). Thus, mechano-protection of the genome by LMNA likely plays a critical role not only in embryonic development, but also in a broad range of adult diseases.

## Statistics

All statistical analyses were performed and graphs were generated using GraphPad Prism 5. All error bars reflect ± SEM. Unless stated otherwise, all comparisons for groups of three or more were analyzed by one-way ANOVA followed by a Dunnett’s multiple comparison test. Pairwise sample comparisons were performed using student’s t-test. Population distribution analyses were performed using the Kolmogorov-Smirnov test, with α = 0.05. Information regarding statistical analyses are included in the figure legends. For all figures, the *p*-values for statistical tests are as follows: n.s. = not significant, \**p*<0.05, **<0.01, ***<0.001 (or #*p*<0.05, ##<0.01 for multiple tests within the same dataset).

## Key resources table

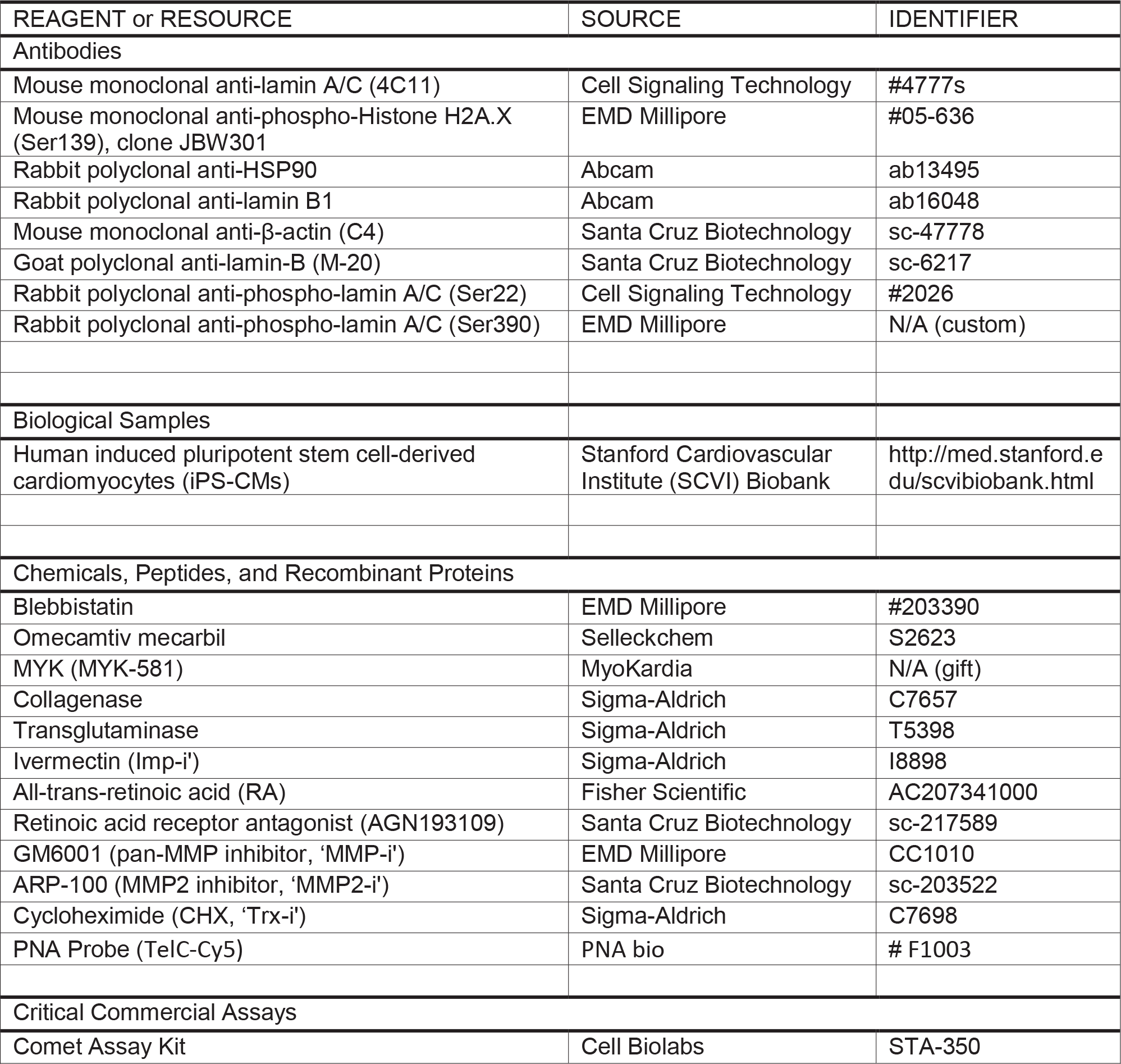

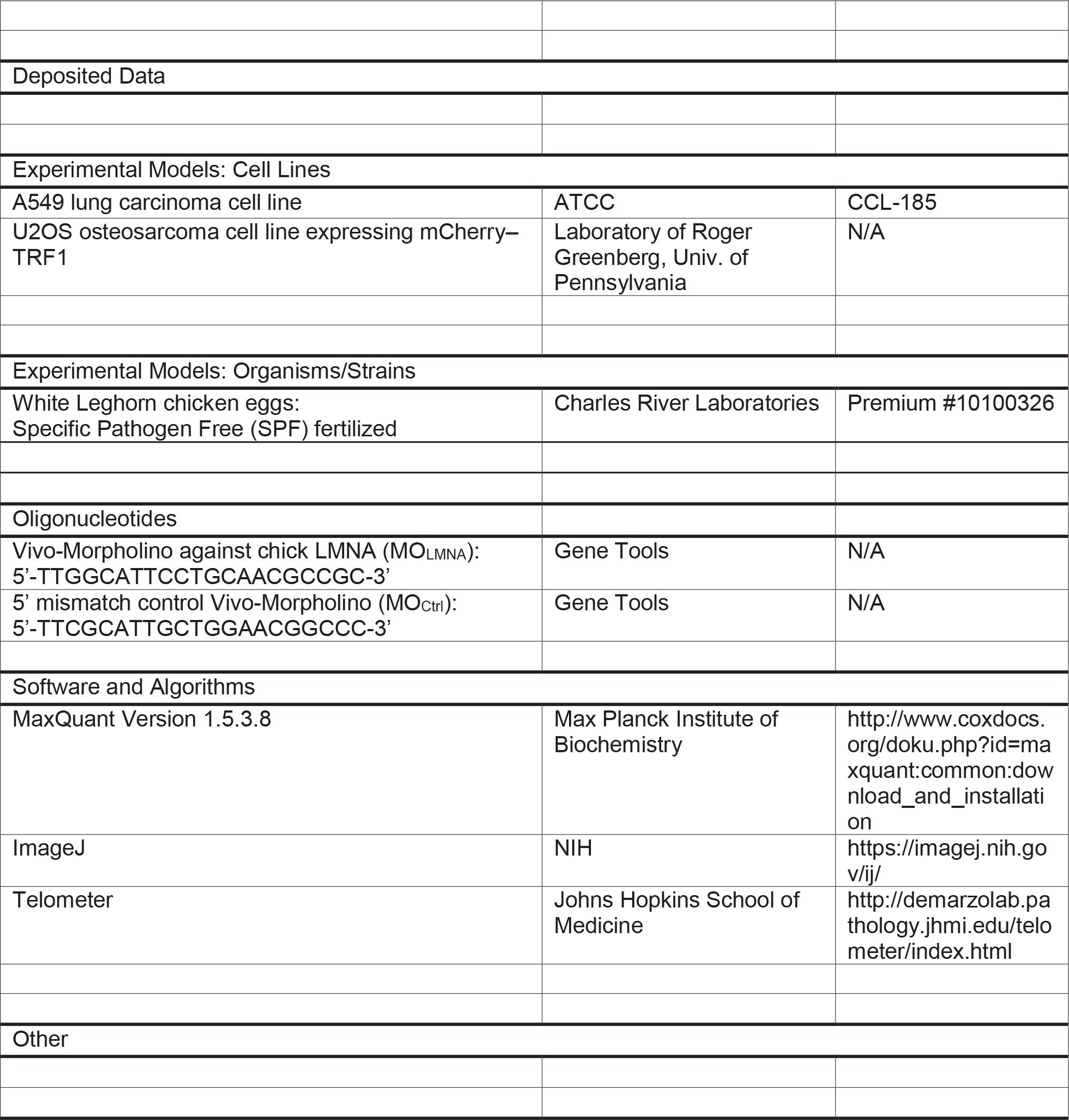

## Acknowledgements

This work was supported by: National Institutes of Health/National Heart Lung and Blood Institute Awards R01 HL124106 and R21 HL128187, National Cancer Institute PSOC Award U54 CA193417, the US–Israel Binational Science Foundation, Charles Kaufman Foundation Award KA2015-79197, and National Science Foundation grant agreement CMMI 15-48571.

## Author Contributions

Conceptualization SC, MV, DED; Investigation SC, MV, AA, SM, KV, II, JI, MT; Validation YX, KZ; Formal Analysis SC, DED; Resources ET, FM, HT, RG, BP; Writing SC, DED; Supervision DED; Funding Acquisition DED.

contractility. Furthermore, arrhythmias and broader conduction defects that are common for many cardiac laminopathies (Fatkin, et al., 1999) provide additional evidence of a potential causal link between DNA damage and the coordinated contractions of CMs (Fig.2). Thus, mechanosensing by LMNA to protect the genome not only plays a critical role during embryonic development, but also has significant clinical implications for a broad range of diseases.

## Supplemental Figure Legends

**Figure S1.**
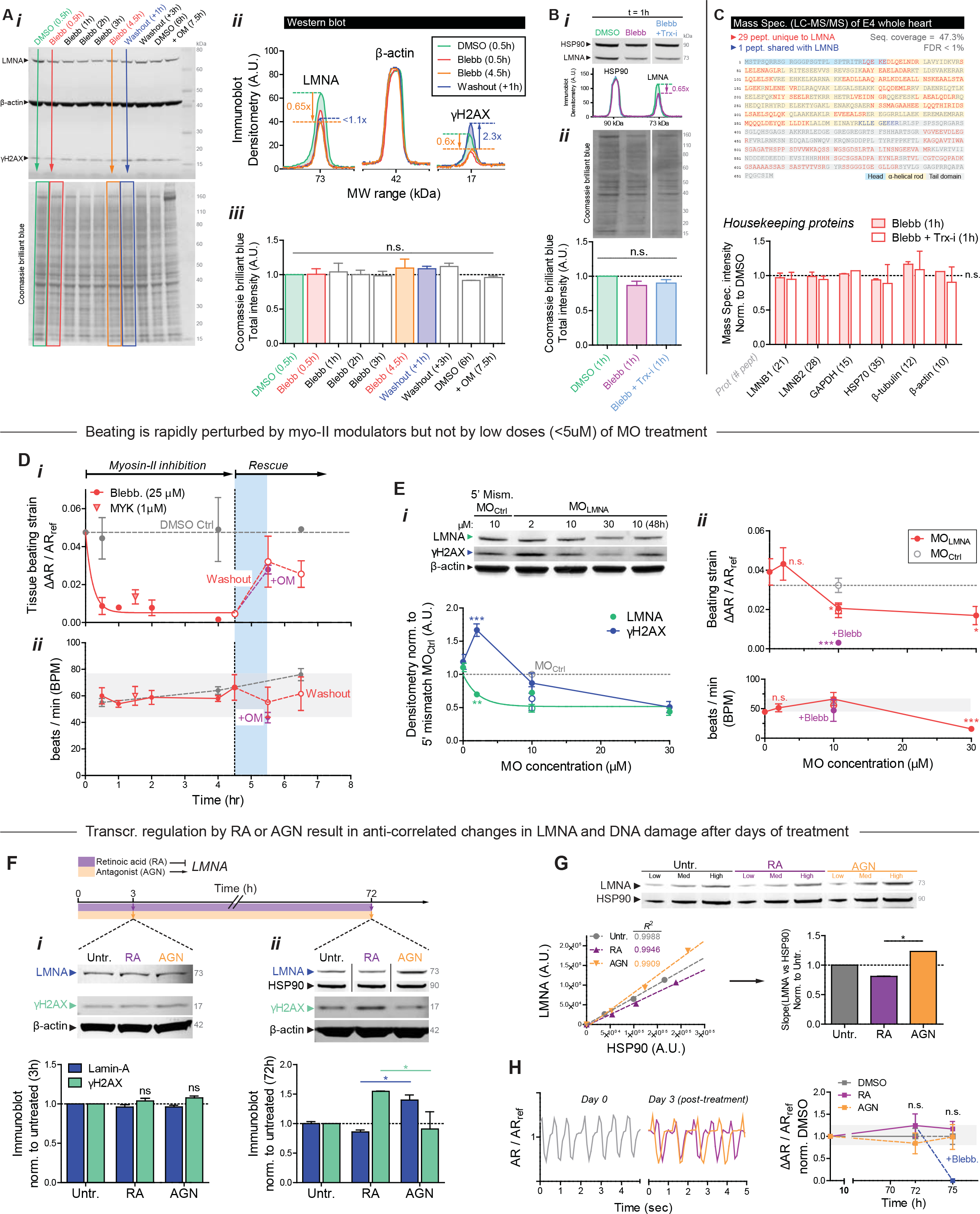
Reversible inhibition of contractility and transcriptional suppression of LMNA result in increased DNA damage, Related to Figures 1 & 2. **(A) (*i*)** Full-length Western blot (top) and corresponding Coomassie Brilliant Blue-stained gel (bottom) for data shown in Fig.1B&C. Key time points are highlighted in colors. **(*ii*)** Densitometry line profiles along the electrophoresis axis (vertical colored arrows) shows significant changes in LMNA and γH2AX, but not β-actin (loading control). **(*iii*)** Coomassie Brilliant Blue total intensity along each lane shows total protein content remains unchanged across conditions. All error bars indicate ±SEM. **(B) (*i*)** WB for a key time point (1h post-blebb treatment) using an alternative housekeeping protein HSP90 as a loading control (top). Densitometry line profiles (bottom) show similar ~35% reduction in LMNA but not HSP90, even in the presence of a protein synthesis inhibitor, (CHX, or ‘Trx-i’). **(*ii*)** Coomassie Brilliant Blue total intensity remains unchanged (<10%) across conditions (n=8 hearts per lysate). **(C)** Mass spectrometry (MS) of heart lysates detects 29 peptides unique to LMNA, and verifies that a broad range of housekeeping proteins, including LMNB1, LMNB2, GAPDH, HSP70, β-tubulin, and β-actin, remain unchanged with blebbistatin treatment (± Trx-i). n=8 hearts per lysate. **(D) (*i*)** Quantification of tissue beating strain ΔAR/AR_ref_ and **(*ii*)** heart rate (beats / min; BPM) in blebbistatin treated hearts. Myosin-II inhibition by blebbistatin rapidly suppresses beating (<30 min), but effects are reversible such that washout of drug with culture medium (± OM) results in near full recovery by 1h. **(E) (*i*)** LMNA and γH2AX immunoblots with different doses of MO. **(*ii*)** Low dowses (<5 µm) have no significant effect on beating strain (ΔAR/AR_ref_) or rate (BPM), but higher doses suppress beating, indicating potential cytotoxicity (one-way ANOVA; *p<0.05, **<0.01). **(F) (*i*)** Retinoic acid (RA) and antagonist to retinoic acid (AGN) treatment in intact embryonic hearts have no observable effect on LMNA levels or DNA damage (γH2AX) at 3h. (n=6 hearts per cond.). **(*ii*)** Significant changes in LMNA and γH2AX are detectable only after 72h treatment, consistent with slow transcriptional modulation. A reduction in LMNA upon RA treatment (72h) is accompanied by an increase in DNA damage as measured by γH2AX, while AGN treatment leads to an upregulation of LMNA coupled to suppression of DNA damage (n=8 hearts per lysate, t-test; \**p*<0.05). **(G)** LMNA WB of RA/AGN treated E4 hearts, with three different loading volumes (Low, Med, High). Bottom left: LMNA vs HSP90 scatter plot (all R^2^ > 0.99); Right: linear fits from LMNA vs HSP90 scatter plot show slopes for AGN > Untr. > RA. **(H)** RA and AGN treatment does not significantly affect contractility of hearts even after 72h of treatment.

**Figure S2.**
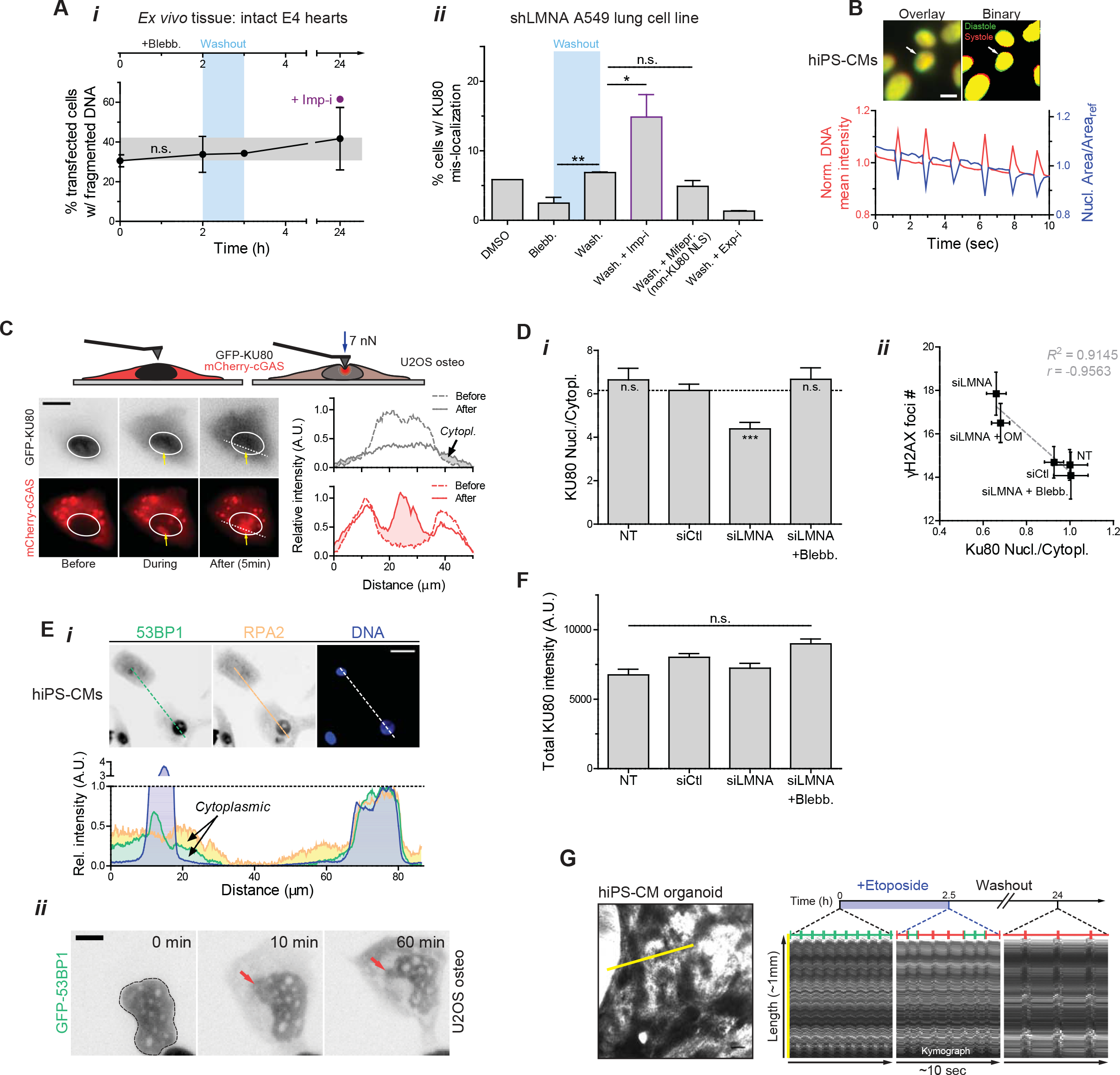
Suppression of LMNA levels in intact embryonic hearts and in beating hiPS-CMs increase rupture under high stress, causing prolonged (>1h) loss of repair factors from the nucleus and accumulation of DNA damage, Related to Figure 3. **(A) (*i*)** Blebbistatin treatment and washout have no significant effect on cell death/viability, as determined by %-transfected cells with fragmented DNA. **(*ii*)** As with embryonic hearts, blebbistatin washout results in an increase in rupture with cytoplasmic mis-localization of KU80. An excess of rupture is seen with washout plus a nuclear import inhibitor (ivermectin, ‘Imp-i’, which inhibits lmpα/β mediated import), but not with washout plus a specific inhibitor of integrase(IN) nuclear import (mifepristone, ‘Mifepr.’) that does not target KU80’s NLS. On the other hand, washout with a nuclear export inhibitor (leptomycin B, ‘Exp-I’) greatly suppresses cytoplasmic mis-localization of KU80 (t-test \**p*<0.05, **<0.01). All error bars indicate ±SEM. **(B)** As seen with nuclei in intact embryonic hearts, ‘nuclear beating’ occurs in hiPS-CMs and can be quantified by changes in nucleus area and DNA mean intensity (condensation/de-condensation of DNA), which are inversely correlated. **(C)** Time-lapse images of nuclei probed with a pointed (<1 µm) Atomic Force Microscopy (AFM) tip (at ~7 nN). Nuclear rupture upon stress is evident in the rapid and stable accumulation of a cytoplasmic protein that binds DNA (GFP-cGAS). **(D) (*i*)** Nuclear/cytoplasmic KU80 IF intensity ratio decreases with siLMNA knockdown, and **(*ii*)** anti-correlates with γH2AX foci count (one-way ANOVA; \*\*\**p*<0.001). **(E) (*i*)** Cytoplasmic mis-localization is not limited to KU80, but applicable to other DNA repair factors including 53BP1 and RPA2 (top: IF images, bottom: intensity profiles along white dashed line). Scale bar = 10 µm. **(*ii*)** Time-lapse images of siLMNA knockdown cells transduced with GFP-53BP1 show that nuclear rupture and cytoplasmic mis-localization occur within minutes and are maintained for at least 1h in culture indicating slow recovery. **(F)** Total (nuclear + cytoplasmic) KU80 abundance is unaffected by siLMNA or blebbistatin treatment. **(G)** Kymograph of beating hiPS-CM organoids generated by tracing length along yellow line with time. Blebbistatin treatment results in reversible inhibition of contractility.

**Figure S3.**
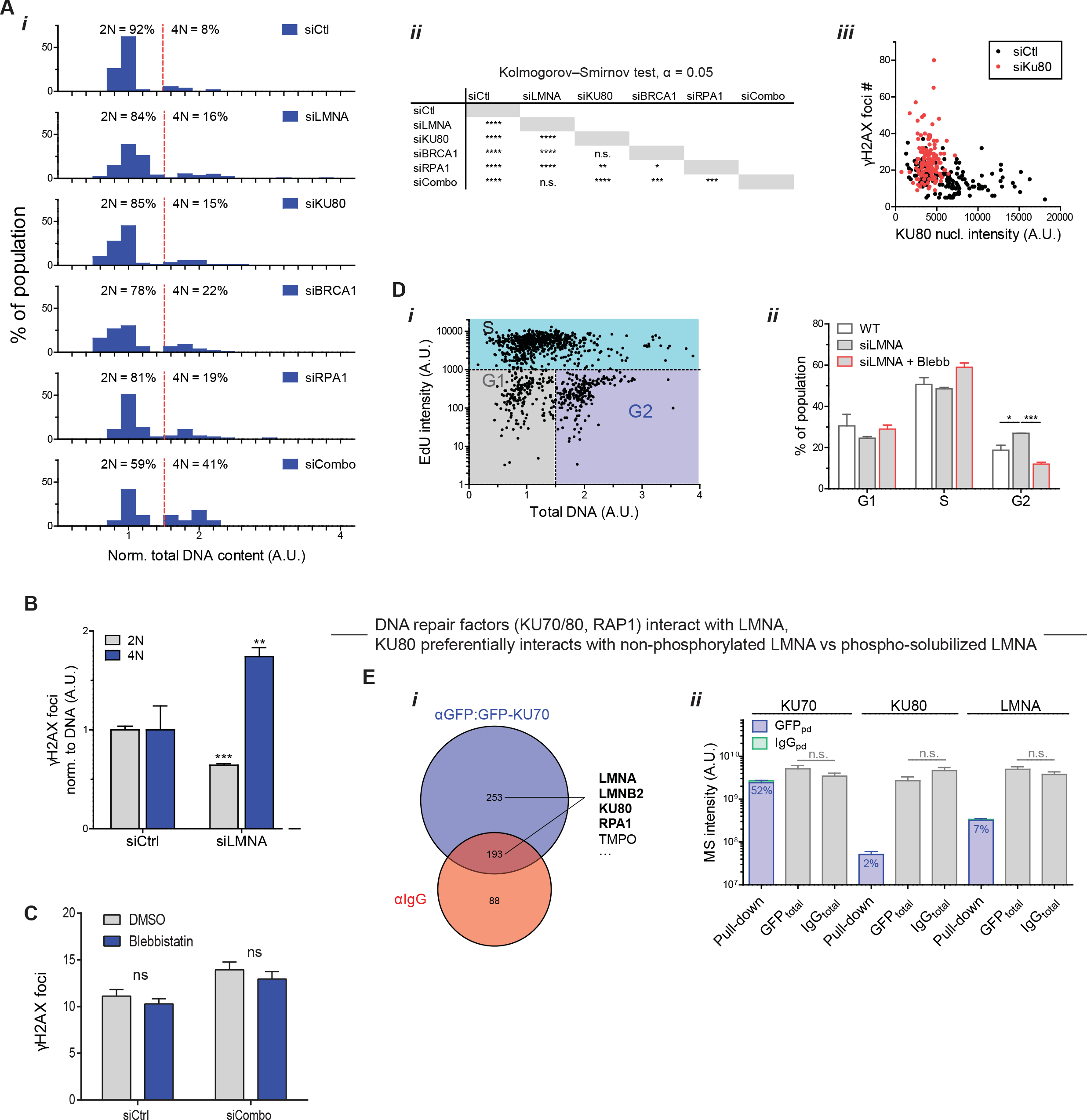
Repair factor loss perturbs cell cycle, but effects are rescued by myosin-II inhibition, Related to Figure 4. **(A) (*i*)** Histograms of normalized DNA content in hiPS-CMs after knockdown of LMNA and DNA repair factors. All repair factor knockdowns results in a higher fraction of ‘4N’ cells, with effects most severe for siCombo. **(*ii*)** Kolmogrov-Smirnov test (α = 0.05) for siRNA treated populations. **(*iii*)** γH2AX foci count inversely correlates with KU80 nuclear IF intensity at the single cell level, consistent with limited repair. All error bars indicate ±SEM. **(B)** DNA damage per total DNA is elevated in 4N cells after LMNA knockdown and decreased in 2N cells, based again on measurements of individual cells. **(C)** Left: Representative scatter plot of EdU intensity versus total DNA content used to determine cell cycle phase (G1, S, and G2). Right: siLMNA treatment increases fraction of cells in G2, but effects are rescued by blebbistatin (t-test \**p*<0.05, ***<0.001). **(D) (*i*)** Venn diagram of nuclear proteins pulled down by co-immunoprecipitation mass spectrometry (CoIP-MS) using anti-GFP on GFP-KU70 expressing cells. **(*ii*)** As expected, KU80 is a strong interaction partner of KU70 (as components of the KU complex), as are lamins-A and B.

**Figure S4.**
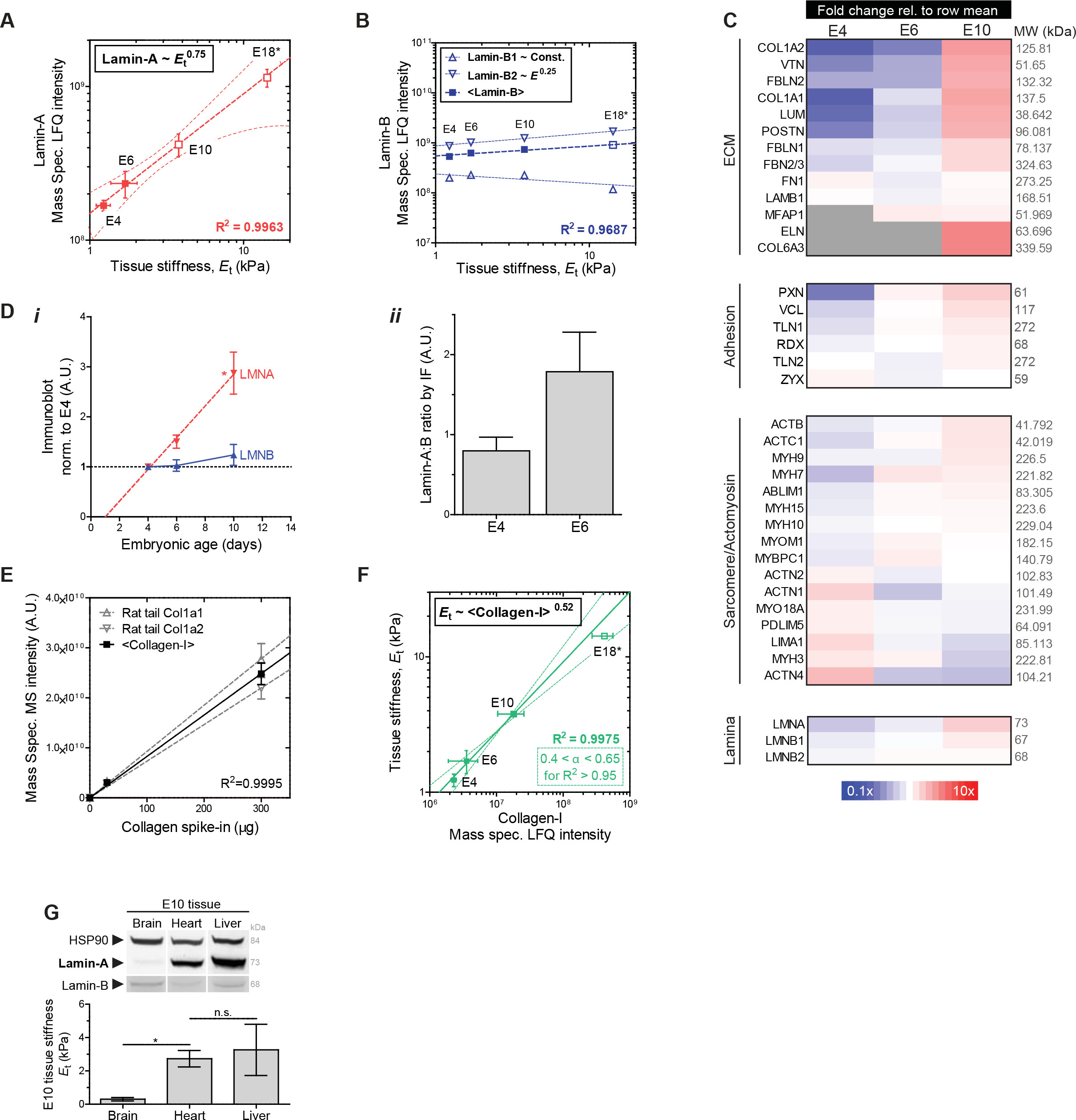
Parallel increases in lamin-A:B, collagen matrix, and tissue stiffness fit a ‘lose it or use it’ model of tension-stabilization of fibers, Related to Figure 5. **(A)** LMNA MS intensity scales with heart tissue stiffness with exponent α ~ 0.75 similar to that for adult tissue proteomes (Swift, et al., 2013). All error bars indicate ±SEM. **(B)** Lamins-B1 & B2 remain comparatively constant throughout development and exhibit much weaker scaling (α < 0.25). **(C)** Heartmap of proteins detected by LC-MS/MS in the ECM, adhesion complexes, sarcomeres, and the nuclear lamina. Proteins were ranked based on the fold-change relative to the average. **(D)** Lamin-A:B ratio quantified by **(*i*)** WB densitometry and **(*ii*)** IF (t-test \**p*<0.05). **(E)** MS measurements of collagen-I calibrated with spike-ins of known amounts of purified collagen-I. Dividing calibrated measurements by reported myocardium volumes (Kim, et al., 2011) gives collagen-I densities far lower than those used in studies of collagen-I gels (Yang, et al., 2009), suggesting lower densities are needed for a stiffness in the kPa range as measured for heart (Fig.4C-*ii*). **(F)** Heart stiffness measurements by micropipette aspiration plotted against collagen-I MS intensity yield power-law scaling comparable to that found for diverse adult tissues (Swift, et al., 2013). **(G)** LMNA & B immunoblots for E10 brain and liver tissue.

**Figure S5.**
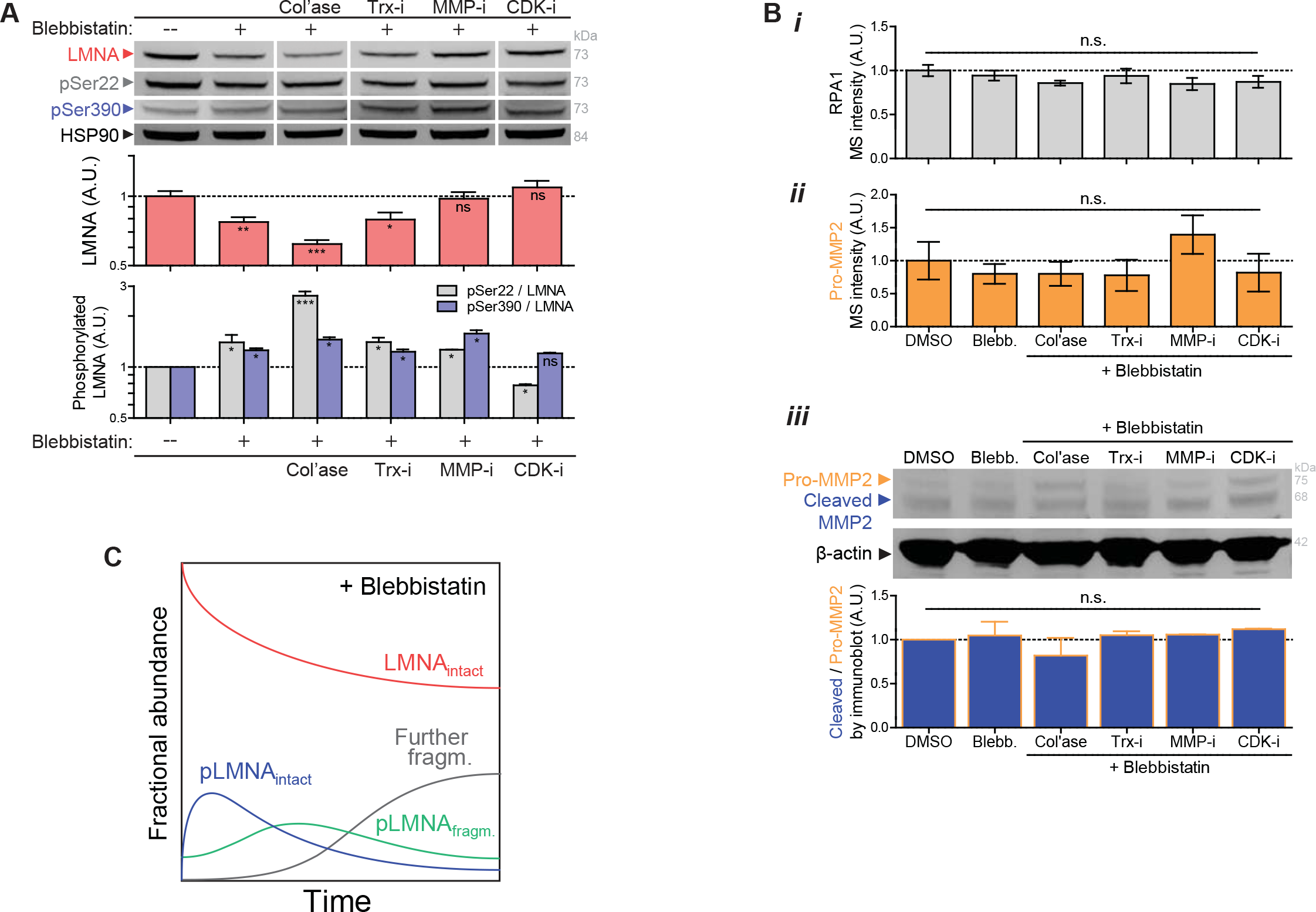
Acute drug perturbations in E4 heart rapidly impact LMNA levels and phosphorylation at multiple sites, but do not affect total abundance of repair factors or MMP2, Related to Figure 5. **(A)** Immunoblot validation of LMNA trends seen in MS profiling of drug perturbations (n>6 hearts per lysate). Immunoblots against phosphorylated Ser22 and Ser390 show that normalized phosphorylation (‘pSer22/LMNA’ and ‘Ser390/LMNA’) increases with blebbistatin-inhibition of actomyosin stress or with collagenase-softening of tissue, but decreases with CDK-i treatment (one-way ANOVA; \**p*<0.05, ** *p*<0.01, \*\*\**p*<0.001). All error bars indicate ±SEM. **(B) (*i*)** RPA1 (DNA repair factor) and **(*ii*)** pro-MMP2 levels are unaffected by drug perturbations (per MS). **(*iii*)** Catalytic activation of MMP2 (‘cleaved/pro-MMP2’ fraction, as measured by immunoblot) is not affected by drug perturbations to contractility and/or collagen matrix. **(C)** Schematic plot of fractional abundance of intact LMNA and its phosphorylated fragments, including intermediates, upon relaxation of stress.

**Figure S6.**
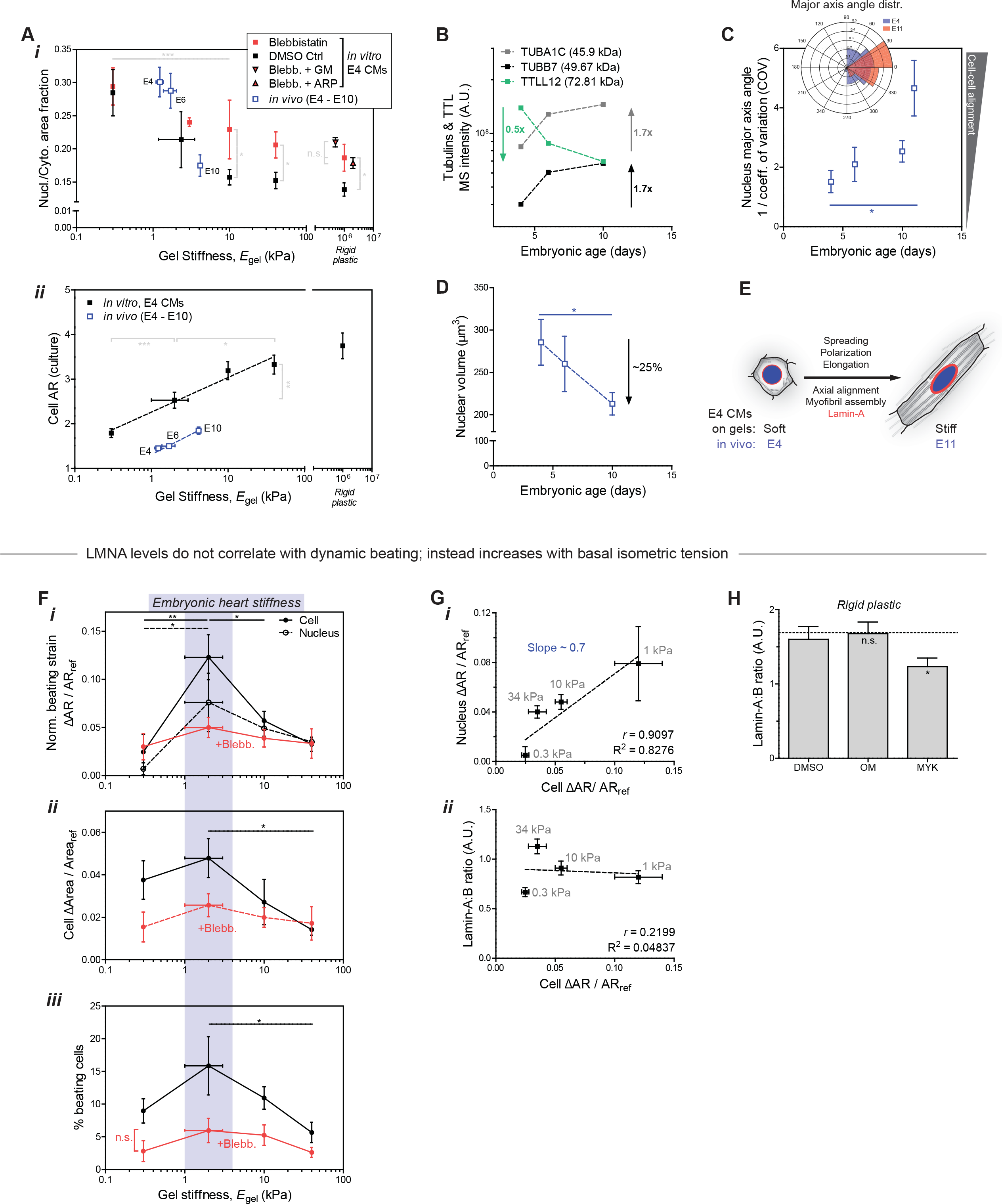
Cell-on-gel morphological trends in CM size, shape, contractility, and LMNA mirror those of *in vivo* hearts without affecting cell cycle/proliferation, Related to Figure 6. **(A) (*i*)** Projected area of the nucleus relative to that of the cell (‘Nucl./Cyto. Area fraction’) decreases **(*ii*)** and cells elongate (increased aspect ratio, AR), as the embryonic heart stiffens and cells undergo hypertrophic growth and spreading from E4 – E10. Isolated E4 CMs cultured on gels likewise exhibit increased cell spreading and elongation on stiffer gels (t-test \**p*<0.05, **<0.01, ***<0.001). All error bars indicate ±SEM. **(B)** Two of the most abundantly expressed α- and β-tubulin isoforms (TUBA1C, TUBB7) increase in level from E4 to E10 and the increase is accompanied by a decrease in TTLL12, which tyrosinates and destabilizes microtubules (Robison, et al., 2016). Trends are consistent with increased stiffness as well as with polarization/elongation of CMs during development (Fig.4D-*ii*, Fig.S6A). **(C)** Axial alignment of cells (anisotropy, quantified as 1/COV of the major axes of nuclei) increases from early (E4) to late (E11) hearts. Upper left inset: major axis angle distribution of E4 and E11 nuclei. **(D)** Nuclear volume (estimated by confocal Z-stack) decreases in development from E4 to E10. **(E)** Varying matrix stiffness alone in E4 CM cultures is sufficient to recapitulate trends in morphology and intracellular organization seen *in vivo* (from E4 to E11). **(F)** Normalized cell and nuclear beating strains measured by **(*i*)** ‘ΔAR/AR_ref_’ and **(*ii*)** ‘ΔArea/Area_ref_’, and **(*iii*)** %-beating cells all exhibit an optimum on gels mimicking the stiffness of embryonic hearts (~2 kPa). Blebbistatin treatment abolishes mechano-sensitivity. (n>10 cells/nuclei per condition). **(G) (*i*)** Nuclear beating strain (‘Nucleus ΔAR/AR_ref_’) correlates well with cell beating strain (‘Cell ΔAR/AR_ref_’), **(*ii*)** but lamin-A:B ratio does not correlate with beating. Lamin-A:B instead couples to average morphology changes in spreading area and elongation that relate to basal isometric tension. **(H)** Lamin-A:B in cells on rigid plastic decreases with MYK inhibition of cardiac myosin contractility (as with blebbistatin treatment), but remains unchanged with OM treatment.

**Figure S7.**
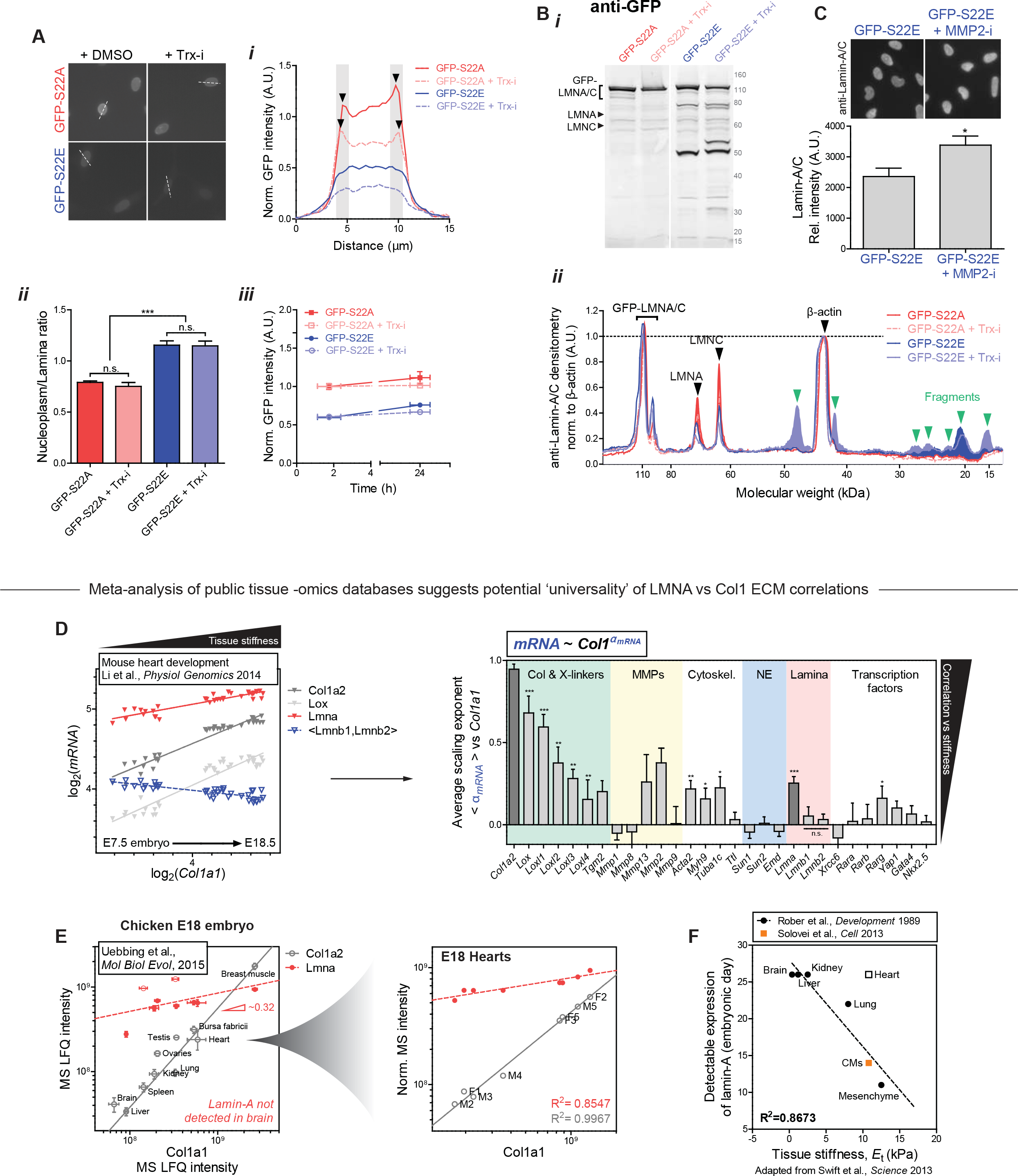
Phosphorylation of LMNA favors degradation by MMP2, Related to Figure 6 & 7. **(A)** Representative images of cells transduced with phsopho-mimetic mutants GFP-S22A and GFP-S22E, with or without protein synthesis inhibitor Trx-i. **(*i*,*ii*)** Line profiles across individual nuclei reveal ‘non-phosphorylatable’ GFP-S22A signal is far more enriched at the lamina than in the nucleoplasm, compared to ‘constitutively phosphorylated’ GFP-S22E (n>15 nuclei). Treatment with Trx-i does not alter nucleoplasm/lamina ratio in either mutant. **(*iii*)** Overall fluorescence intensity of GFP-S22A is ~50% higher than that of GFP-S22E. Trx-i induces a minor ~10% decrease in either case. All error bars indicate ±SEM. **(B)** Immunoblots with anti-lamin-A/C and anti-GFP reveal multiple low-MW degradation fragment bands (green triangles) in the GFP-S22E mutant which are absent in the S22A mutant. **(*ii*)** Line intensity profile of anti-lamin-A/C immunoblot (*i*, left). Low-MW degradation fragments that are present in the GFP-S22E mutant but not in the GFP-S22A mutant, are shaded in blue, with green arrows indicating distinct bands. **(C)** GFP fluorescence intensity of GFP-S22E expressing cells increases upon MMP2-I (ARP-100) treatment, consistent with inhibition of degradation. **(D)** Representative transcriptomics dataset for mouse embryonic hearts obtained from NCBI’s Gene Expression Omnibus (GEO) Database. Log-log plot shows normalized mRNA expression vs *Col1a1*. As expected for obligate heterotrimer subunits of collagen-I, *Col1a2* correlates robustly with *Col1a1*, with scaling exponent (= slope on a log-log plot), α_*Col1a2*_ ~ 1. Lmna also increases with *Col1a1*, although with slightly weaker scaling α_*Lmna*_ ~ 0.3, indicating potential feedback to gene expression. Right: bar graph of average scaling exponents (vs *Col1a1*) for proteins of interest in the ECM, cytoskeleton, nuclear envelope, as well as several transcription factors implicated in mechanosensing and/or cardiac development. **(E)** Proteomics dataset for diverse E18 chick embryonic tissues, adapted from Uebbing et al. (Uebbing et al., 2015). Lmna again increases with Col1a1&2, with exponent aLmna ~ 0.3. Right inset: datapoints for heart samples plotted separately reveal similar scaling. **(F)** Tissue-dependent timing of initial LMNA expression in early embryos correlates well with the stiffness that a given tissue eventually achieves in adult (adapted from (Swift, et al., 2013; Rober, et al., 1989)).

